# Premature thymic involution in young *Foxn1^lacz^* mutant mice causes peripheral T cell phenotypes similar to aging-induced immunosenescence

**DOI:** 10.1101/2024.04.23.590170

**Authors:** Shiyun Xiao, Seung Woo Kang, Kimberly E. Oliva, Wen Zhang, Kimberly D. Klonowski, Nancy R. Manley

## Abstract

The thymus is a primary lymphoid organ generating self-restricted and self-tolerant naïve T-cells. Early in life the thymus starts to involute, resulting in decreased naïve T-cell output which may be more self-reactive, leading to an increased prevalence of autoimmunity. A decrease in the transcription factor FOXN1 is an early event in thymic involution. Using the *Foxn1^lacz^* model, we studied how premature thymic involution affects the thymic microenvironment, thymocytes, and peripheral T cell immunity. We found that early thymic involution led to aged-like thymic epithelial cells that resulted in aged-like thymocyte phenotypes, with a significant decrease in CD4+ single-positive T-cells. We also observed severe lymphopenia in *Foxn1^lacz^* mice caused by the premature decrease in T-cell production, resulting in a peripheral T-cell phenotype similar to *de novo* aged peripheral T-cells. Moreover, following T cell receptor stimulation, *Foxn1^lacz^* peripheral T cells had reduced IL-2 secretion and strong initial IFN-g responses, resembling aged wild-type peripheral T-cell responses. Lastly, influenza response in *Foxn1^lacz^* had a reduction in some aspects of T cell responses to influenza infection, similar to thymectomized mice. Our study shows an independent and direct impact of premature thymic involution on both thymopoiesis and peripheral immune niches likely contributing to immunosenescence and inflammaging as observed in the elderly population.

## Introduction

Thymus size and function peak early in life and then start to decline with age, defined as thymic involution. As a result, naive T cell (T_N_) export declines and T cell receptor (TCR) diversity and self-tolerance is impaired over time (Bains et al. 2009; Shanley et al. 2009). These changes are regulated by thymic epithelial cell (TEC)-based mechanisms and crosstalk between TECs, non-TEC stroma, and developing thymocytes (Chinn et al. 2012). While hematopoietic stem cells (HSC) that seed the thymus functionally decline with age, thymic involution occurs prior to this decline and is stromal-intrinsic (Mackall et al. 1998; Zhu et al. 2007).

Reduced expression of Forkhead Box N1 (FOXN1) is an early event in thymic involution (Chen et al. 2009; Ortman et al. 2002). As a key regulator of TEC biology throughout life (Chen et al. 2009; Manley & Condie 2010; Nowell et al. 2011; Blackburn et al. 1996; Nehls et al. 1996; Zook et al. 2011; Su et al. 2003), over-expression of FOXN1 can preserve TECs and delay thymic involution (Zook et al. 2011; Bredenkamp et al. 2014). *Foxn1* is incredibly dosage-sensitive, and small decreases cause changes in TEC differentiation and function (Chen et al. 2009; Nowell et al. 2011; Bredenkamp et al. 2014). Although *Foxn1* expression in the thymus is restricted to TECs, it indirectly affects the maturation of thymic vasculature and mesenchyme (Nowell et al. 2011; Bryson et al. 2011), orchestrating nearly all aspects of thymic function and maintenance.

Our previous report of the *Foxn1^lacz^* mouse model showed that *Foxn1* expression in TECs is normal in fetal and early postnatal stages but declines after postnatal day 7 to about 40% of wild type levels, causing a progressive reduction in thymopoiesis and premature thymic involution (Chen et al. 2009). Since down-regulation of *Foxn1* expression occurs similarly in TECs , preceding aging-related involution (Chen et al. 2009), *Foxn1^lacz^* models age-related thymic involution in young mice and allows investigation of how thymic involution contributes directly to peripheral immunosenescence.

In this study, we used single-cell RNA sequencing and flow cytometry to dissect the independent effect of premature thymic involution on thymopoiesis and peripheral immune function. Using *Foxn1^lacz^* mice, we show that an early decrease in *Foxn1* leads to an aged TEC phenotype in young animals, resulting in *de novo* aged-like thymopoiesis. The premature decline in thymic output resulted in lymphopenia in *Foxn1^lacz/lacz^* mice and formed an aged wild type-like peripheral immune system and *in vitro* T-cell responses suggesting immunosenescence and inflammaging in young *Foxn1^lacz/lacz^*. Together, our findings provide insight into the direct impact of premature thymic involution on peripheral T-cells independent from hematopoietic stem cells or physiological aging.

## Results

### Single-cell RNA sequencing of CD45-total thymic stroma shows an aged TEC phenotype in 1-month-old *Foxn1^lacz/lacz^*

In order to study how the relative level of *Foxn1* affects the thymic stroma transcriptome at a cellular level, we submitted CD45-thymic stroma of 1-month-old *Foxn1^+/+^* (+/+) males and *Foxn1^lacz/lacz^* (Z/Z) males for single-cell RNA sequencing. We merged these datasets with a batch correction to a published 18-months-old C57BL6/J wild type (wt) female dataset (Kousa et al. 2023) to compare *de novo* aged thymus to thymus with low *Foxn1* expression. We detected all previously described stromal subsets based on their signature marker expressions. We identified 10 TEC clusters [*Epcam,* MHC class II], 4 endothelial clusters [*Pecam1*, *Cdh5*], 3 fibroblast clusters [*Pdgfra, Col1a1*], 2 pericyte clusters [*Pdgfrb, Acta2, Myl9*], mesothelial cells [*Upk3b, Nkain4*], and rare non-myelinating Schwann cells [*Gfap, Ngfr, S100b*] (Kousa et al. 2023; Hu et al. 2018) (Fig. 3.1 A).

**Figure 3.1.**
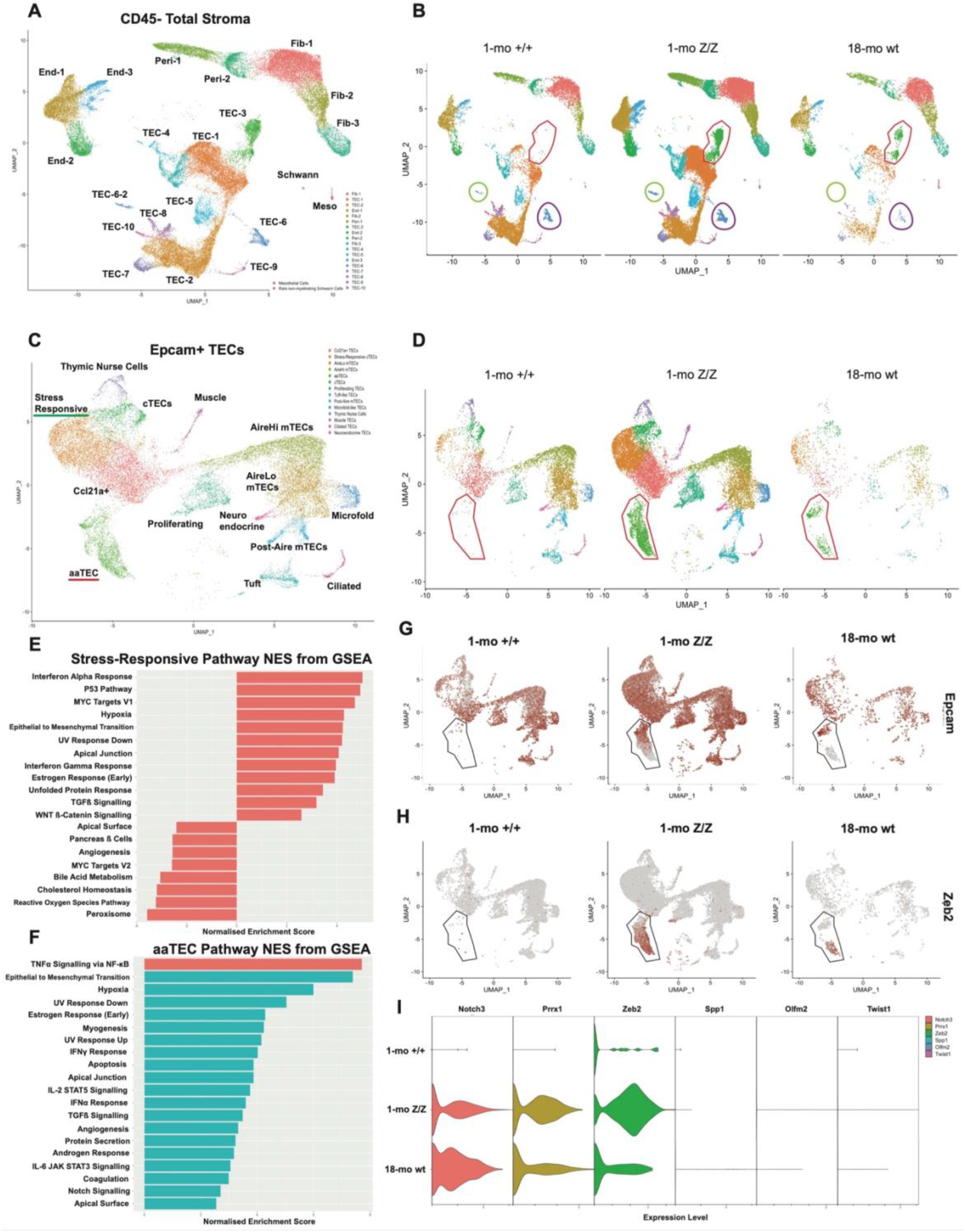
scRNAseq analysis of CD45-total thymic stroma and thymic epithelial cells revealed an aged TEC phenotype in 1-month-old Foxn1*^lacz^*. (A) UMAP plot visualization of the CD45-total stroma. (B) UMAP plot visualization of the CD45-total stroma split by age/genotype. Red circles indicate TEC-3 that were only largely present in 1-month-old (mo) Foxn1 Z/Z and 18-mo wt. Lime green circles indicate TEC-6-2 and purple circles indicate TEC-6 that were absent in 18-mo wt. (C) UMAP plot visualization of the Epcam+ TECs. (D) UMAP plot visualization of the Epcam+ TECs split by age/genotype. Red circles indicate aaTEC that were only present in 1-mo Foxn1 Z/Z and 18-mo wt. (E) Top 20 hallmark pathways ranked by normalized enrichment score for stress-responsive cTECs vs. rest of the TECs. (F) Top 20 hallmark pathways ranked by normalized enrichment score for aaTECs vs. rest of the TECs. (G) Feature plot of an epithelial cell marker, *Epcam*, split by age/genotype. (H) Feature plot of a mesenchymal cell marker, *Zeb2*, split by age/genotype. (I) Violin plot of an aaTEC cluster, split by age/genotype. Abbreviations: *mo,* months old; *wt*, wild type; *Z/Z, Foxn1^lacz/lacz^*.

Major differences were observed in TECs between the three datasets. Two TEC populations (TEC-6-2, TEC-9) were absent in the 18-months-old wt sample compared to 1-month-old +/+ and Z/Z (Fig. 3.1 A, B). A subset of TEC-6 and TEC-9 were identified as muscle TECs [*Myog, Myl1, Actc1*] and ciliated TECs [*Spag8, Dnah12, Dynlrb2*] (Michelson et al. 2022), respectively. Of note, ciliated TECs express high levels of *Cldn3*, *Cldn4*, and *Plet1* (Depreter et al. 2008), suggesting a decrease in TEC progenitor cell population in the 18-months-old wt). We also observed a TEC population (TEC-3) that was scarce in 1-month-old +/+ samples but was highly enriched in both 1-month-old Z/Z & 18-month-old wt (Fig. 3.1 B).

We subsampled *Epcam*+ MHC Class II genes+ clusters to acquire a better resolution of TEC populations (Fig. 3.1 C). We identified 12 previously defined TEC populations (Brennecke et al. 2015; Miragaia et al. 2018; Bornstein et al. 2018; Baran-Gale et al. 2020; Park et al. 2020; Bautista et al. 2021; Michelson et al. 2022) and two unknown TEC populations (Fig. 3.1 D). To identify these new TEC populations, we used differentially expressed gene analysis (DEG) and gene set enrichment analysis (GSEA). GSEA of one of the unique cTEC population revealed IFN*α* response, P53 pathway, hypoxia, and epithelial to mesenchymal (EMT) transition, suggesting it was stress-responsive (Fig. 3.1 E). GSEA of the other unique TEC population revealed TNF*α* signaling via NF-*κ*B and EMT. DEG analysis showed that this population was previously described as aging-associated TECs (aaTECs) (Kousa et al. 2023) (Fig. 3.1 F). Concordant to Kousa et al. (2023), the aaTEC population is undergoing EMT based on their loss of epithelial markers (*Epcam, Pax1*) and acquisition of mesenchymal markers (*Zeb2, Notch3, Prrx1*), which we confirmed (Fig. 3.1 G-I). However, aaTECs in 1-month-old Z/Z had a slightly different transcriptome compared to 18-month-old wt. One-month-old Z/Z aaTECs had 1.6 times higher *Zeb2* expression but 16.3 times less *Spp1* compared to 18-month-old wt aaTECs. aaTECs comprised 10.4% of 1-month-old Z/Z TECs, whereas this increased to 22.4% of 18-month-old wt TECs. Overall, our single-cell RNA sequencing data suggests a decrease in *Foxn1* resulted in the premature occurrence of stress-responsive cTECs and aaTECs, resembling aged TEC phenotypes.

### Premature thymic involution alters thymic selection and accelerates CD4 and CD8 SP thymocyte maturation in *Foxn1^lacz^* mutant mice

Down-regulation of *Foxn1* expression in TECs from postnatal day 7 causes premature thymic involution in young adult *Foxn1^lacz^* mice with a significant reduction of medullary thymic epithelial cells (mTECs), early thymic progenitors (ETPs), and total thymocyte number (Chen et al. 2009; Xiao et al. 2018). These changes are similar to the profiles of age-related thymic involution in naturally aging wt mice (Min et al. 2004). To identify differences in thymocyte developmental deficiency between premature thymic involution of young (2-months-old) *Z/Z* mice and age-related thymic involution of aged (18-months-old) wt mice, we compared their thymocyte phenotype to young (2-months-old) *Foxn1^+/lacz^* control mice (+/Z). The percentages of both CD4 and CD8 SP thymocytes were significantly reduced in young Z/Z mutants, but in old wt mice, only CD8 SP thymocytes were significantly reduced compared to +/Z (Fig. 3.2 A). To determine the effects of thymic positive selection on the production of both CD4 and CD8 SP thymocytes, we measured the expression of positive selection initiative molecules CD69, CD5, and TCRb (Žuklys et al. 2016; Saini et al. 2010). Compared to +/Z control, the expression of CD69 (Fig. 3.2 B) and CD5 (Fig. 3.2 C) on TCRβ^int^ DP thymocytes and TCRβ^hi^ SP were reduced in both young Z/Z mutants and old wt mice, indicating decreased thymic positive selection.

**Figure 3.2.**
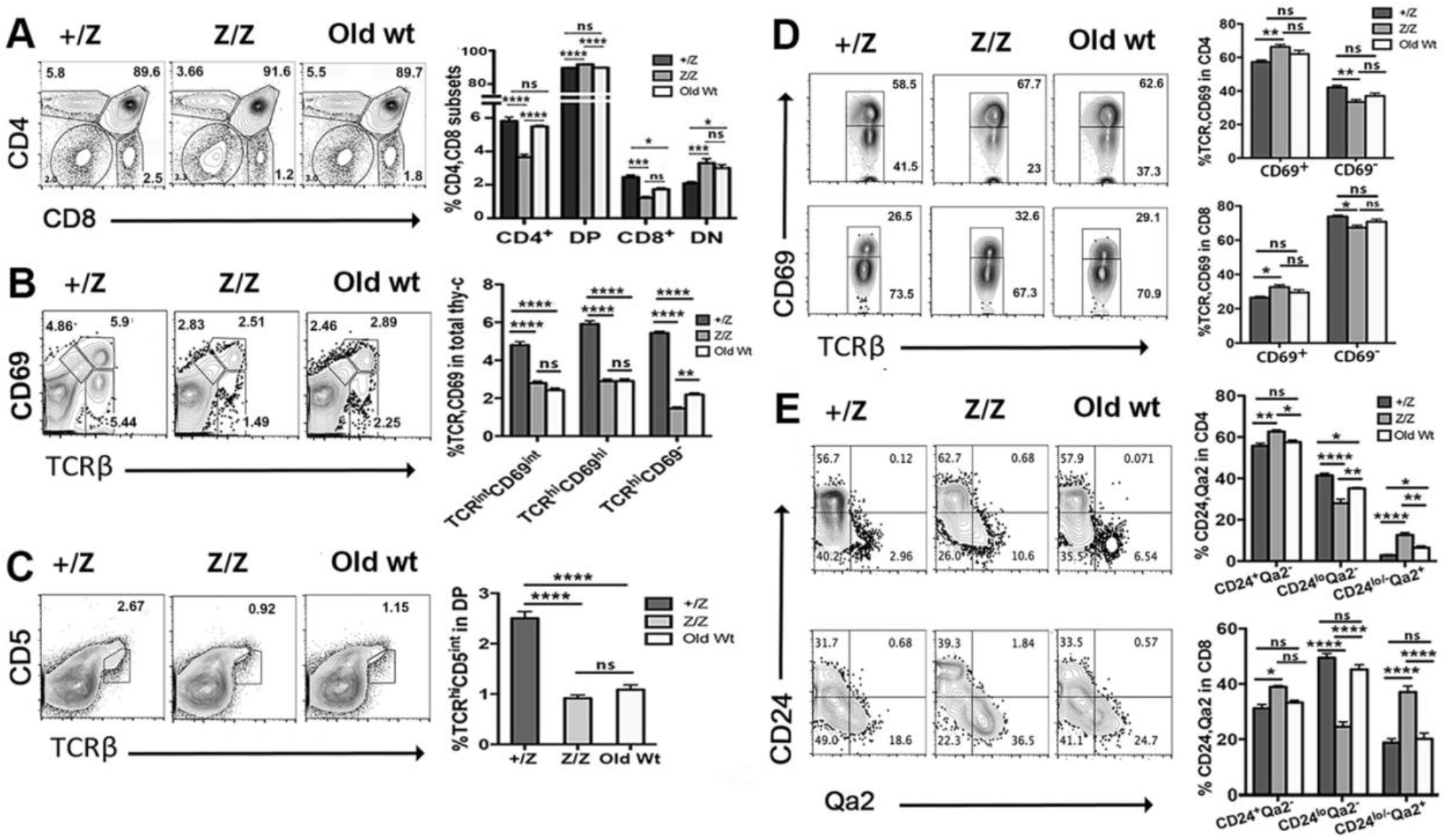
Thymic selection and maturation of CD4 and CD8 SP thymocytes in young *Foxn1^lacz^* mutants. Flow cytometry analysis of thymocytes from +/Z (2-months-old), Z/Z (2-months-old) and old wt mice (18-months-old). **(A).** Expression of CD4 and CD8 on total thymocytes. Percentage of DN, DP, CD4, and CD8 subsets on the right. **(B).** Expression of CD69 and TCRβ on total thymocytes. Percentage of TCRβ^int^CD69^int^, TCRβ^hi^CD69^hi^ and TRCβ^hi^CD69^-^ subsets on the right. **(C).** Expression of CD5 on DP thymocytes. Percentage of TCRβ^hi^CD5^int^ on the right. **(D).** Expression of CD69 and TCRβ on gated CD4 (top) and CD8 (bottom) SP thymocytes. Percentages of TCRβ^+^CD69^+^ and TCR^+^βCD69^-^ on the right. **(E).** Expression of CD24 and Qa2 on gated CD4 (top) and CD8 (bottom) SP thymocytes. Percentage of CD24^+^Qa2^-^, CD24^lo^Qa2^-^ and CD24^lo/-^Qa2^+^ on the right. (A, B, D and E): Two-way ANOVA; (C): One-way ANOVA. P values: ****≤0.0001, ***≤0.001, **≤0.01, *≤0.05.

We measured the negative selection process by analyzing the down-regulation of CD69, CD24, and CCR7 on post-selection of TCRβ^hi^ CD4 and CD8 SP thymocytes (Žuklys et al. 2016; Ueno et al. 2004). The percentages of CD69^-^ TCRβ^hi^ (Fig. 3.2 D) T cells were significantly reduced in young Z/Z but not in old wt mice. However, CD24^lo^Qa2^-^ CD4 but not CD8 (Fig. 3.2 E), CD24^lo^CCR7^+^CD4, and CD24^lo^CCR7^lo^CD8 (S-Fig. 3.1 A) T cells were significantly reduced in both Z/Z and old wt mice. These results indicated that negative selection was increased in both young Z/Z and old wt mice with some differences. After negative selection, CD4 and CD8 SP thymocytes stay in the thymic medulla for approximately 15 days for further function maturation, with one of the important changes being the up-regulation of Qa2 expression (Vernachio et al. 1989). The percentage of CD24^lo/-^Qa2^+^ cells was significantly increased in both CD4 and CD8 SP thymocytes in young Z/Z mutants and only mildly increased in old wt mice compared to +/Z controls (Fig. 3.2 E). A BrdU incorporation test showed that the proliferation potential of Qa2^+^ cells was similar between mouse genotypes (S-Fig. 3.1B), suggesting that the increase in CD24^lo/-^ Qa2^+^ CD4 and CD8 SP thymocytes in Z/Z mutants was likely due to accelerated function maturation rather than increased proliferation. Together, young Z/Z mutants and old wt mice showed similarly reduced thymic positive selection and maturation (Fig. 3.2 B, C) but differences in thymic negative selection, with young Z/Z mutants showing increased negative selection (Fig. 3.2 D, E, S-Fig. 3.1 A).

### The proliferation of developing thymocytes is altered in young *Foxn1^lacz^* mutant mice

The size of the postnatal thymus is controlled by the expansion of early-stage thymocytes at CD4^-^CD8^-^ DN and CD4^+^CD8^+^ DP stages. DN proliferation is limited by the availability of stromal niches, but DP proliferation is limited intrinsically (Prockop & Petrie 2004). IL-7 and c-KitL are two key cytokines supporting the proliferation of early thymocytes at the DN stage (Peschon et al. 1994; von Freeden-Jeffry et al. 1995). We measured the receptors of IL-7 and c-KitL on DN thymocytes to determine if alteration of cytokine-mediated DN proliferation could be the cause of the reduced number of total thymocytes. Compared to +/Z control (1.5-months-old), both young Z/Z (1.5-months-old) and old wt (18-months-old) showed a significant increase of IL-7 receptor (IL-7Rα / CD127) expression on DN1 subsets, whereas only young Z/Z mutants showed an increase on DN3 subsets. (Fig. 3.3A, B). c-Kit expression on DN1 and DN2 subsets was significantly reduced in young Z/Z and old wt (Fig. 3.3C-E). Additionally, there was a reduction in DN2 proliferation but an increase in DN3 proliferation in young Z/Z compared to controls (Fig. 3.3F, G). Old wt DN thymocytes displayed no changes in proliferation compared to controls (Fig. 3.3F, G). The proliferation capabilities of thymocytes in later stages were also significantly increased in DN, DP, CD4, and CD8 subsets in young Z/Z but not in old wt thymocytes (Fig. 3.3H, I). These results suggest while premature thymic involution in *Foxn1^lacz^* altered thymocyte proliferation, it is not likely the reason behind the reduction in total cellularity of the Z/Z thymus.

**Figure 3.3.**
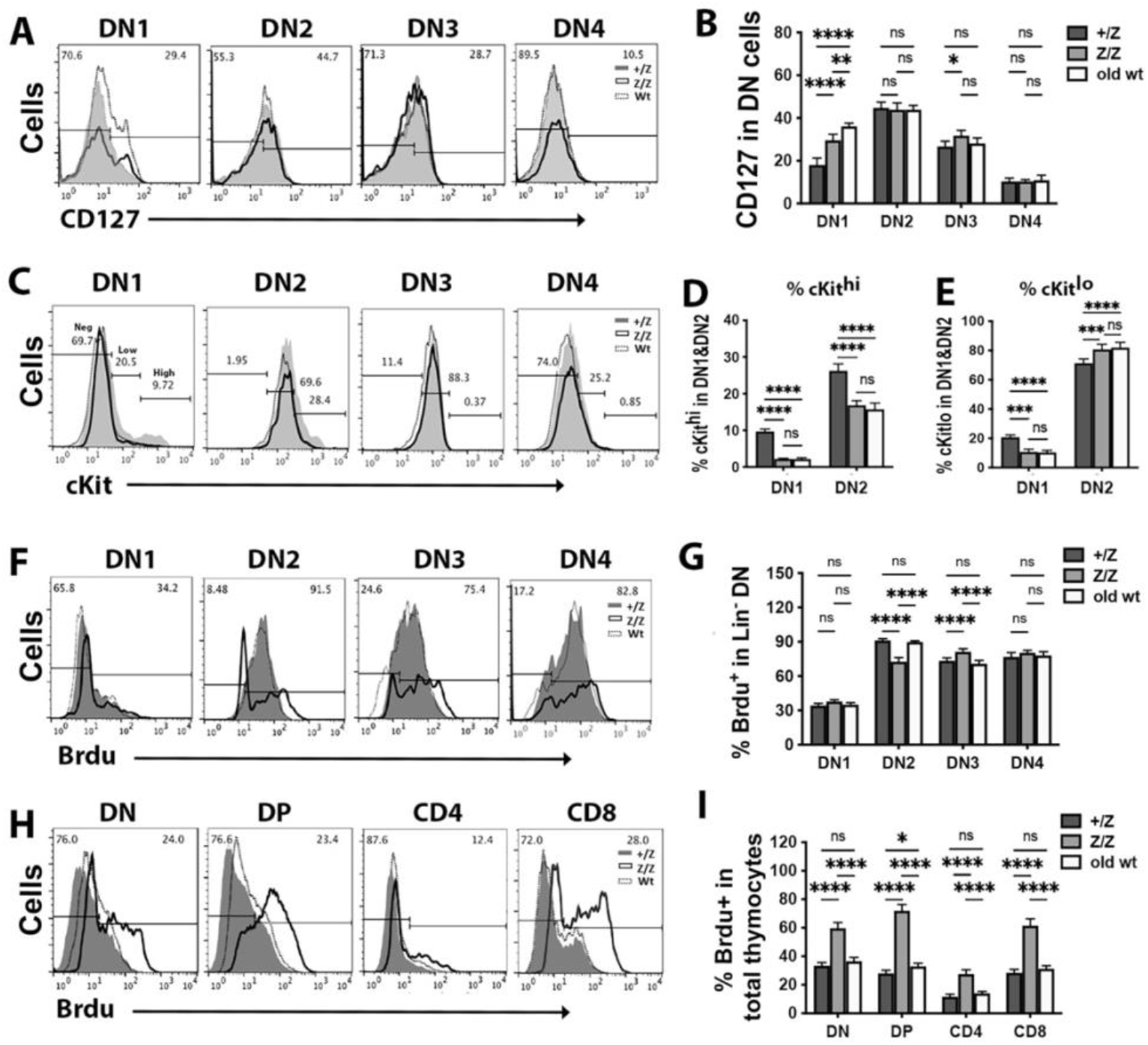
The proliferative capability of thymocytes was slightly increased in young *Foxn1^lacz^* mutants similar to aged wild type mice. **(A-I).** Flow cytometric analysis of thymocytes from +/Z (1.5-months-old), Z/Z (1.5-months-old), and old wt (18-months-old) mice. **(A).** Histogram of CD127 expression on subsets of DN thymocytes. **(B).** Summary data of percentage of CD127 on DN1-4 subsets. **(C).** Histogram of cKit expression on subsets of DN thymocytes. **(D).** Percentage of cKit^hi^ on DN1 and DN2 subsets. **(E).** Percentage of cKit^lo^ on DN1 and DN2 subsets. **(F).** Histogram of Brdu staining on subsets of DN thymocytes. **(G).** Summary data of percentage of Brdu^+^ cells on DN1-4 subsets. **(H).** Histogram of Brdu staining on DN, DP, CD4, and CD8 thymocytes. **(I).** Summary data of percentage of Brdu^+^ cells in DN, DP, CD4, and CD8 thymocytes. Two-way ANOVA. P values: ****≤0.0001, ***≤0.001, **≤0.01, *≤0.05.

We further tested the effect of thymic involution on thymocytes by measuring apoptosis with annexin V staining. Apoptotic cells increased in DN1, DN2, CD4 SP, and CD8 SP stages (Fig. 3.4A, B) in both young Z/Z and old wt mice compared to +/Z control. Together, these results indicate that although there are some differences in proliferation and apoptosis in thymocyte developmental stages of young Z/Z and old wt mice, they are quite similar at DN1 and DN2 stages. Apopototic profile changes causing the reduction of ETPs could be a key factor contributing to thymocyte reduction in these mice (Xiao et al. 2018; Min et al. 2004).

**Figure 3.4.**
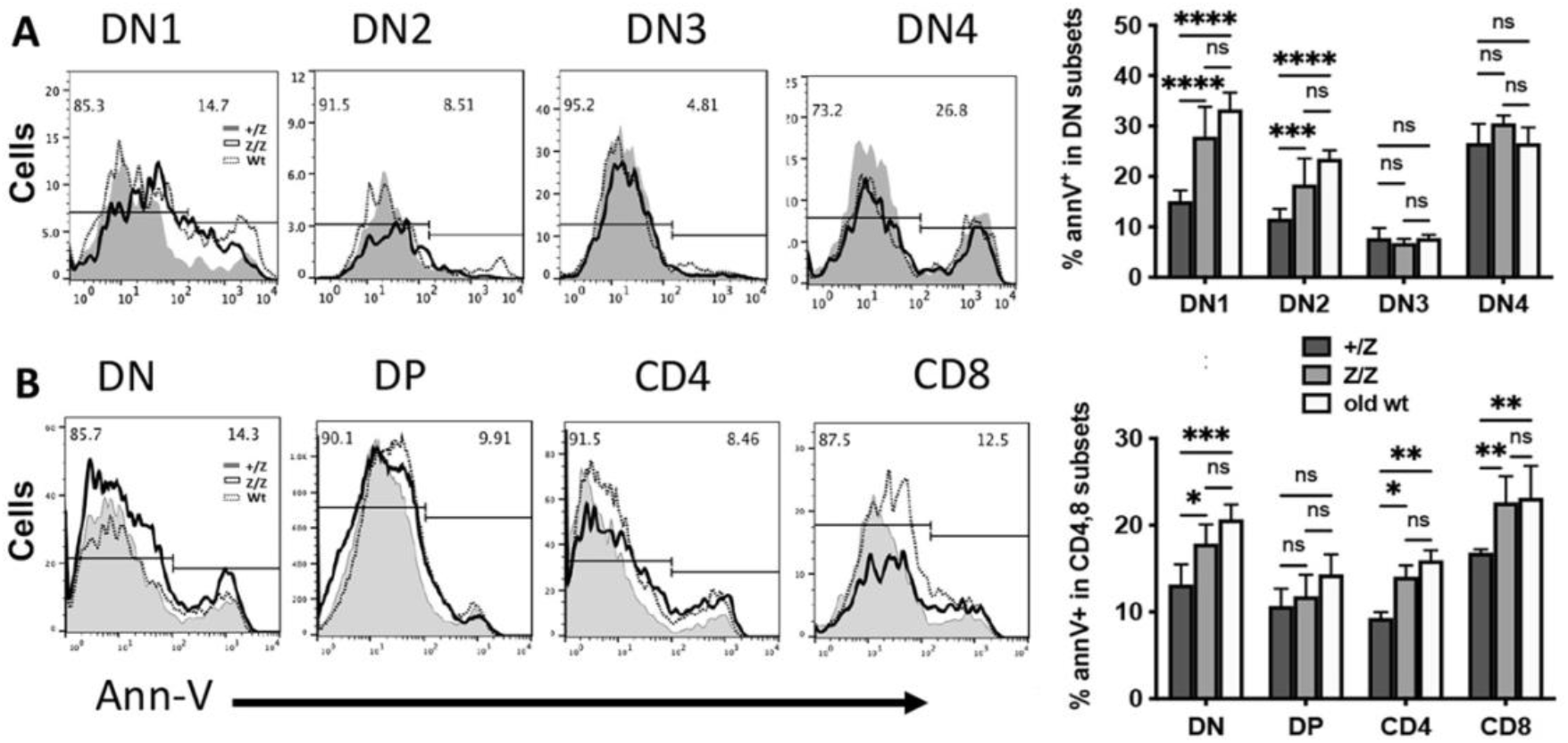
Premature thymic involution in young *Foxn1^lacz^* mice caused an increase in thymocyte apoptosis similar to aged wild type mice. Flow cytometric analysis of thymocytes from +/Z (1-month-old), Z/Z (1-month-old), and old wt (15-months-old) mice by Annexin V staining. **(A).** Histogram of annexin V expression on subsets of DN thymocytes. Summary data of percentage of annexin V^+^ cells in DN1-4 subsets. **(B).** Histogram of annexin V expression on subsets of DN, DP, CD4, and CD8 thymocytes. Summary data of percentage of annexin V^+^ cells in DN, DP, CD4, and CD8 thymocytes. (A and B): Two-way ANOVA. P values: ****≤0.0001, ***≤0.001, **≤0.01, *≤0.05.

### Naïve CD4 and CD8 T-cells are decreased in young *Foxn1^lacZ^* spleen

Since decreased *Foxn1* resulted in aging-like TEC phenotypes, we wanted to investigate how the peripheral immune niches were affected. We submitted total peripheral mononuclear cells (PMCs) from spleen of 1-month-old +/+ and Z/Z mutants with 18-months-old C57BL6/J wt for single-cell RNA sequencing. We detected 7 follicular B-cell clusters [*Cd19, Ighd, Cr2, Cd22*], 1 plasma cell/plasmablast cluster [*Jchain, Iglv1, Derl3, Eaf2*], 4 T-cell clusters [*Cd3e, Cd3g, Cd4, Cd8a*], 1 NK cluster [*Ccl5, Klra4, Klrd1*], 1 cDC cluster [*Clec9a, Xcr1, Clec10a*], 1 pDC cluster [*Siglech, Irf7, Irf8*], 1 monocyte cluster [*Fn1, Clec4a1, Sirpb1c*], 1 basophil cluster [*Prss34, Fcer1a, Cpa3, Mpo*], 1 macrophage cluster [*Ms4a2, Ccl9, Cyp11a1*], and 2 neutrophil clusters [*Wfdc21, Ltf, Ngp, Lcn2*] (Schulze et al. 2022) (Fig. 3.5 A). We observed a decrease in a follicular B-cell population accompanying an increase in plasma cell/plasmablast cluster in 18-month-old wt samples compared to both 1-month old +/+ and Z/Z (Fig. 3.5 B). We also observed an increase in both CD4 and CD8 T cell populations in the 18-month-old wt sample compared to +/+ and Z/Z (Fig. 3.5 B).

**Figure 3.5.**
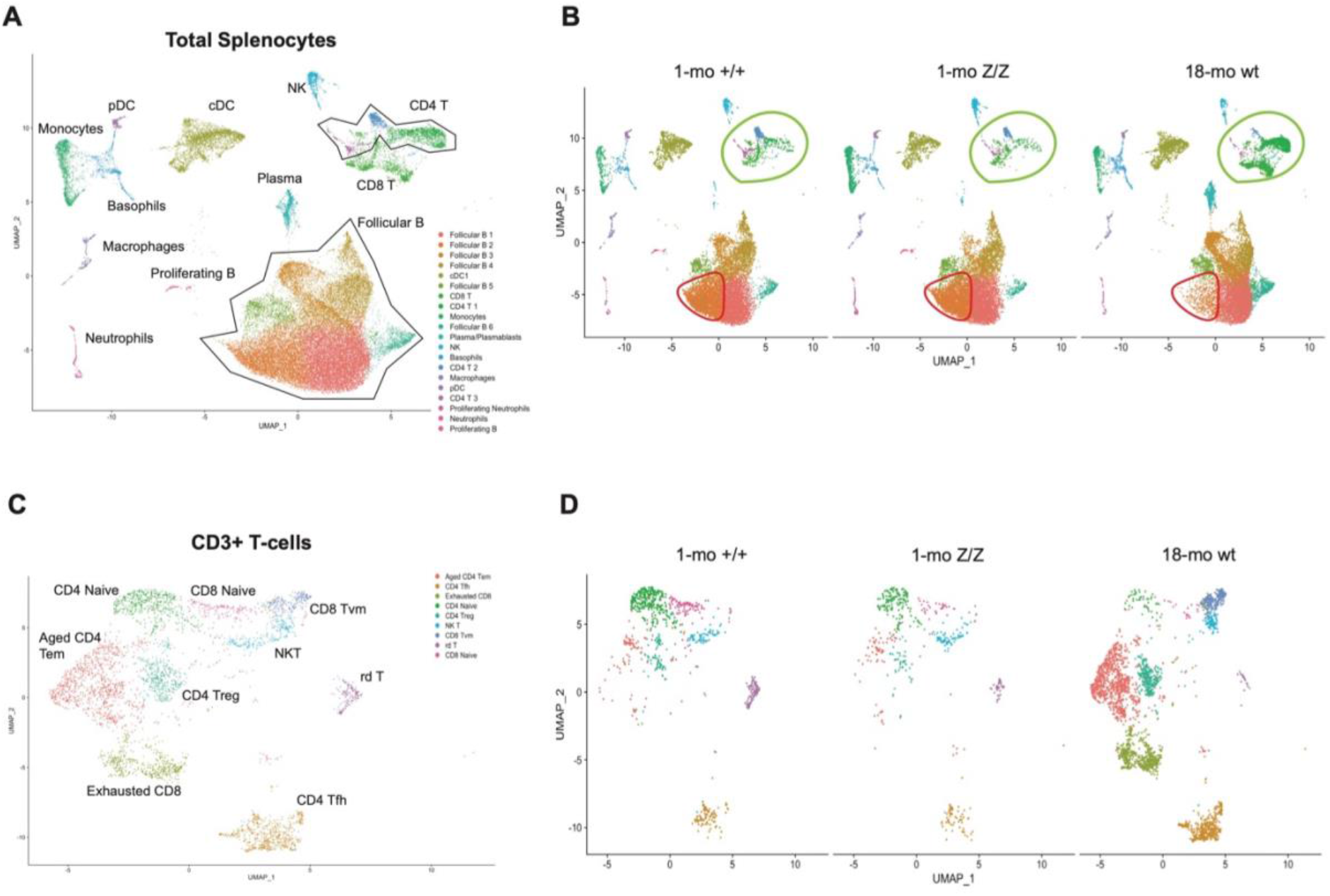
Total splenocyte scRNAseq comparison showed decreased naïve CD4 & CD8 populations in young *Foxn1^lacz^* mice. (A) UMAP plot visualization of total splenocytes. (B) UMAP plot visualization of total splenocytes split by age/genotype. Red circles indicate a follicular B-cell cluster that is exclusively decreased in 18-mo wt. Lime circles indicate CD4 and CD8 single-positive (SP) T-cells. (C) UMAP plot visualization of CD3+ T-cells. (D) UMAP plot visualization of CD3+ T-cells split by age/genotype. Abbreviations: *mo*, months old; *Z/Z*, *lacZ*; *WT*, wild type.

We subsampled *Cd3*+ clusters to have a higher resolution in T-cell subset changes (Fig. 3.5 C). The 18-months-old wt sample showed unique T-cell clusters, such as CD8 virtual memory-like T cells [*Cd8a, Cd44^Hi^, Cd49d^Lo^*] (Hussain & Quinn 2019), aged CD4 T cells [*Cd4, Tnfrsf4k, Tnfsf8, Ctla4, Nt5e*] (Yang et al. 2020), and CD8 exhausted T cells [*Cd8a, Pdcd1, Lag3, Havcr2, Prdm1*] (Jiang et al. 2015), reflecting their age and exposures during their lifetime (Fig. 3.5 D). To understand the direct effect of decreased *Foxn1* in the thymus on peripheral T cell changes, we compared 1-month-old +/+ and Z/Z. There was only a slight decrease in the proportion of CD4 naïve T cells in Z/Z, 35.6% in +/+ compared to 31.9% in Z/Z. Conversely, there was approximately a 45% decrease in the proportion of CD8 naïve T cells in Z/Z, 12.1% in +/+ compared to 6.8% in Z/Z. Additionally, there was a substantial decrease in the proportion of γδ T-cells in Z/Z and old wt. 12.9% of CD3+ T-cells were γδ T in WT, 8.4% in ZZ, and 0.9% in old wt. Overall, these data suggest that the aging TEC phenotype in 1-month-old Z/Z led to decreased splenic naïve CD4 and CD8 T-cell populations compared to +/+ control, similar to 18-month-old wt.

### Peripheral lymphocytes from young *Foxn1^lacz^* mice show an age-related phenotypic change comparable to aged wild type mice

To further evaluate the effects of premature thymic involution on peripheral lymphocytes in young *Foxn1^lacz^*, we used flow cytometry to compare the phenotypic changes of splenic lymphocytes between +/Z controls (2.5-months-old), Z/Z mutants (2.5-months-old), and old wt mice (18-months-old). The percentage of CD4 T cells was significantly reduced in young Z/Z and slightly reduced in old wt mice, but no significant changes were observed in CD8 cells compared to +/Z controls (Fig. 3.6 A). However, the expression CD4 and CD8 were both downregulated on SP T cells from young Z/Z and old wt mice (Fig. 3.6 B, C). Resembling an aged wt lymphocyte profile, T cells from young Z/Z showed significantly increased effector memory T cells (T_em_) and a reduction of naïve T cells (T_n_) (Fig. 3.6 D). Consistent with this, young Z/Z and old wt mice had similar trends in the percentages of TCRβ+ cells and B cells (TCD19+) compared to +/Z controls (S-Fig. 3.2 A-C). The percentage of natural killer (NK) cells and granulocytes in the spleen were only increased in young Z/Z (S-Fig. 3.2 D-G), but the percentage of Treg cells had no significant difference between mice (data not shown).

**Figure 3.6.**
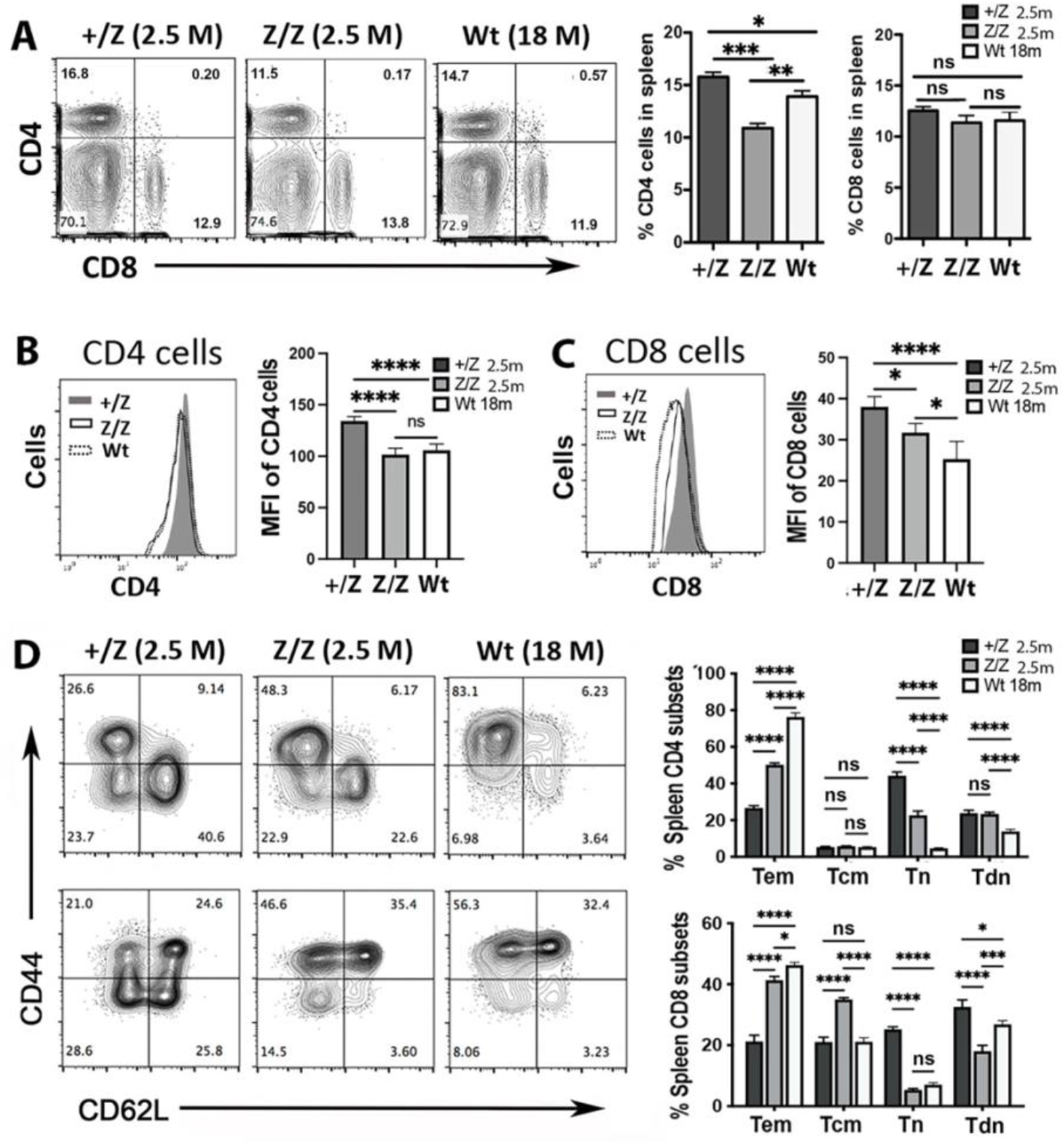
Young *Foxn1^lacz^* mice showed age-related phenotypic changes in peripheral lymphocytes. Flow cytometric analysis of spleen cells from +/Z (2.5-months-old), Z/Z (2.5 months-old), and aged wt (18-months-old) mice. **(A).** CD4- and CD8-expressing spleen cells. Right, percentage of CD4 and CD8 T cells of total spleen cells. **(B).** Histogram of CD4 fluorescence density on gated CD4 T cells. CD4 mean fluorescence density (MFI) is shown to the right. **(C).** Histogram of CD8 fluorescence density on gated CD8 T cells. CD8 mean fluorescence density (MFI) is shown to the right. **(D).** Expression of CD44 and CD62l on gated CD4 (top) and CD8 (bottom) spleen T cells. Percentages of effector memory (Tem), central memory (Tcm), naïve (Tn), and double negative (DN) (CD62L^-^CD44^-^) subsets of T cells are to the right. (A-C): One-way ANOVA; (D): Two-way ANOVA. P values: ****≤0.0001, ***≤0.001, **≤0.01, *≤0.05. Abbreviations: *M,* months.

### Young *Foxn1^lacz^* mice show mild peripheral lymphopenia analogous to aged wild type mice

In wt mice, increased thymocyte production enlarges the pool of peripheral lymphocytes during ontogeny, which remains stable even with thymic involution (Surh & Sprent 2005; Mackall et al. 1997). However, in *Foxn1^lacz^* mice, the production of thymocytes dramatically decreases on postnatal day 7, prematurely affecting the pool of peripheral lymphocytes (Chen et al. 2009). To measure the size of the peripheral lymphocyte pool, we collected cells from the spleen and lymph nodes (LNs) of 2-3-months-old +/Z and Z/Z and 15-20-months-old wt mice. The total number of peripheral mononuclear cells (PMC) was significantly reduced in young Z/Z mutants and old wt mice compared to +/Z control (Fig. 3.7 A). The total number of CD4 and CD8 T cells was also reduced in young Z/Z and old wt mice (Fig. 3.7 B, C). For both CD4 & CD8 T-cells in Z/Z and old wt, the most reduced subset was naïve T cells, and effector memory T cells (T_em_) were the most abundant, especially in old wt mice (Fig 3.7 D, E). To test the peripheral lymphopenia status in *Foxn1^lacz^* mice, we transferred CFSE-labeled CD45.1^+^ CD4 and CD8 T cells into young +/Z, Z/Z, and old wt (CD45.2^+^) mice (S-Fig. 3.3 A). Although most CD45.1^+^ donor cells remained CFSE^hi^ in the host, there was a slight reduction in the percentage of CFSE^hi^ cells in young Z/Z and old wt mice compared to the +/Z mice (S-Fig 3.3 B-F). However, the total number of CD45.1^+^ donor cells recovered from young Z/Z and old wt after transfer was significantly reduced compared to +/Z control (S-Fig. 3.3 G). These results suggest that both young Z/Z and old wt mice show a slight peripheral lymphopenia status that drives homeostatic proliferation of lymphocytes; however, the peripheral environment does not support the survival of proliferated cells. Consistent with lymphopenia, proliferating CD4 and CD8 T cells increased (Fig. 3.7 F, G) and showed similar T-cell subset profiles in young Z/Z and old wt with the exception of CD8 T_em_ subset (Fig. 3.7 H, I), which are uniquely elevated in old wt mice. Also, the percentage of apoptotic cells was increased in both CD4 and CD8 T cells in young Z/Z and old wt mice (Fig. 3.7 J, K). Combined, these results indicate that peripheral lymphopenia in young Z/Z mutants and old wt mice led to homeostatic lymphocyte proliferation, and a concurrent increase in apoptosis significantly reduced the total T-cell number compared to +/Z controls.

**Figure 3.7.**
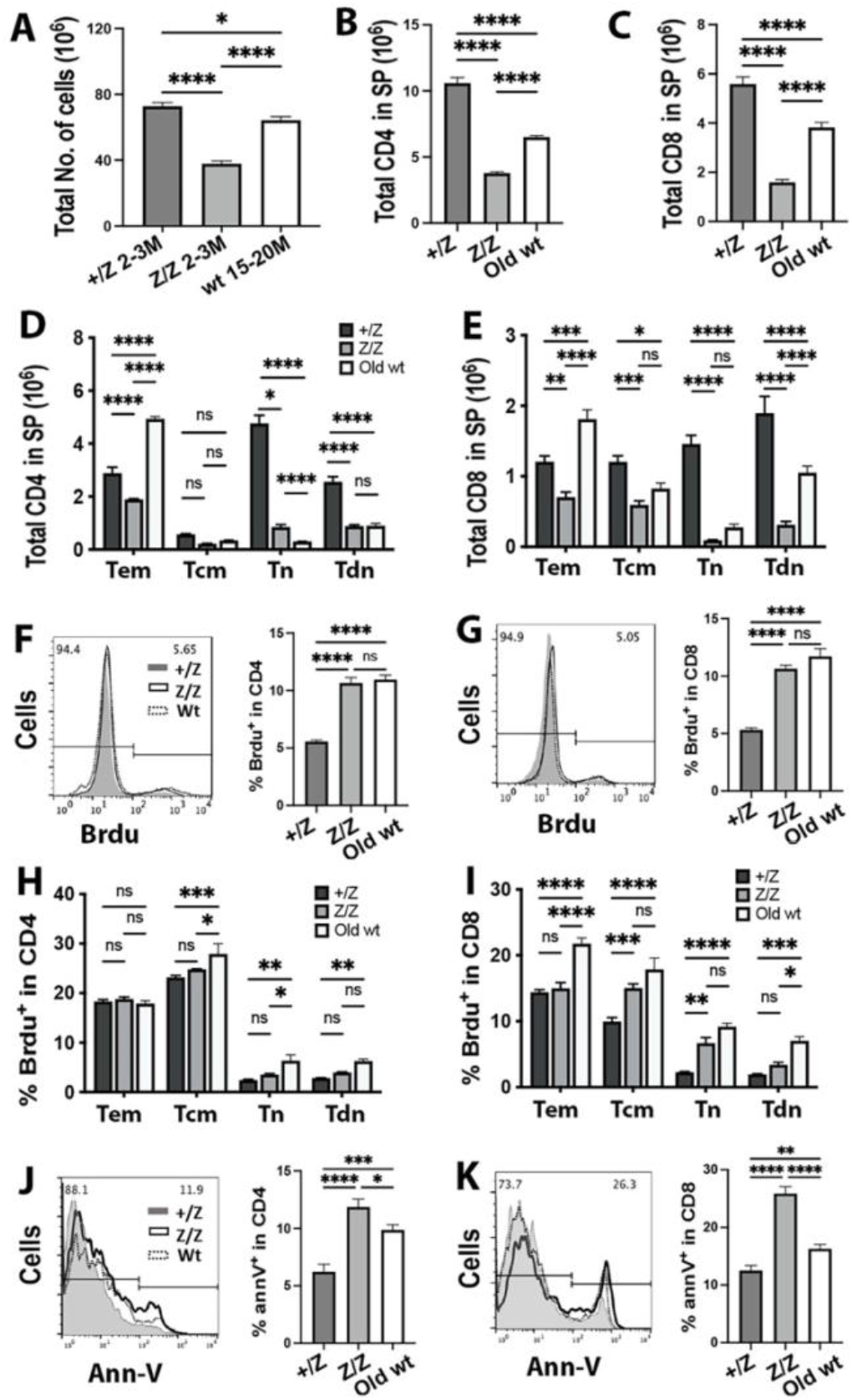
Young *Foxn1^lacz^* mice displayed mild peripheral lymphopenia. **(A).** Histogram of the total number of peripheral lymphocytes prepared from spleen and lymph nodes (LNs) of +/Z (2-3 months-old), Z/Z (2-3 months-old), and old wt (15-21 months-old) mice. **(B and C).** Histogram of the total number of CD4 T cells (B) and CD8 T cells (C) in spleen. **(D and E).** Histogram of the total number of CD4 T cells subsets (D) and CD8 T cell subsets (E) in spleen. **(F and G).** Histogram of Brdu staining CD4 T cells (F) and CD8 T cells (G). Percentages of Brdu^+^ CD4 and CD8 T cells shown to the right. **(H and I).** Histogram of Brdu^+^ CD4 T cell subsets (H) and CD8 T cell subsets (I). **(J and K).** Histogram of Annexin V staining CD4 T cells (J) and CD8 T cells (K). Percentages of Annexin V^+^ CD4 and CD8 T cells shown to the right. (B-K): Cells analyzed in graphs were prepared from +/Z (2 mos), Z/Z (2 mos), and old wt (18 mos) mice. (A-C, J and K): One-way ANOVA; (D, E, H, and I): Two-way ANOVA. P values: ****≤0.0001, ***≤0.001, **≤0.01, *≤0.05. Abbreviations: Tem, Effector memory T-cells; Tcm, Central memory T-cells; Tn; Naïve T-cells.

### T cells from young *Foxn1^lacz^* and aged mice display weak TCR signaling and impaired proliferation

Since young Z/Z mutants and old wt displayed lymphopenia, with a smaller peripheral lymphocyte pool than +/Z control, we analyzed the proliferative capability of these cells after TCR stimulation. To do this, we isolated splenic and LN-derived CD4 T cells and co-cultured them with or without aCD3+CD28 antibodies *in vitro*. Compared to +/Z mice, CD4 T cells from Z/Z and old wt had ∼ 50% reduction in proliferation after stimulation (Fig. 3.8 A). Phenotypic analysis of co-cultured cells 48 hours after activation showed that both activation markers CD69 and CD25 were significantly reduced on CD4 T cells from both Z/Z and old wt compared to +/Z (S-Fig. 3.4 A-E). These results suggest that CD3-TCR signal-induced activation was impaired in both young Z/Z mutants and old wt mice, which is consistent with the down-regulation of TCRβ expression in CD4 and CD8 T cells from both Z/Z and old wt mice (Fig. 3.8 B, C).

**Figure 3.8.**
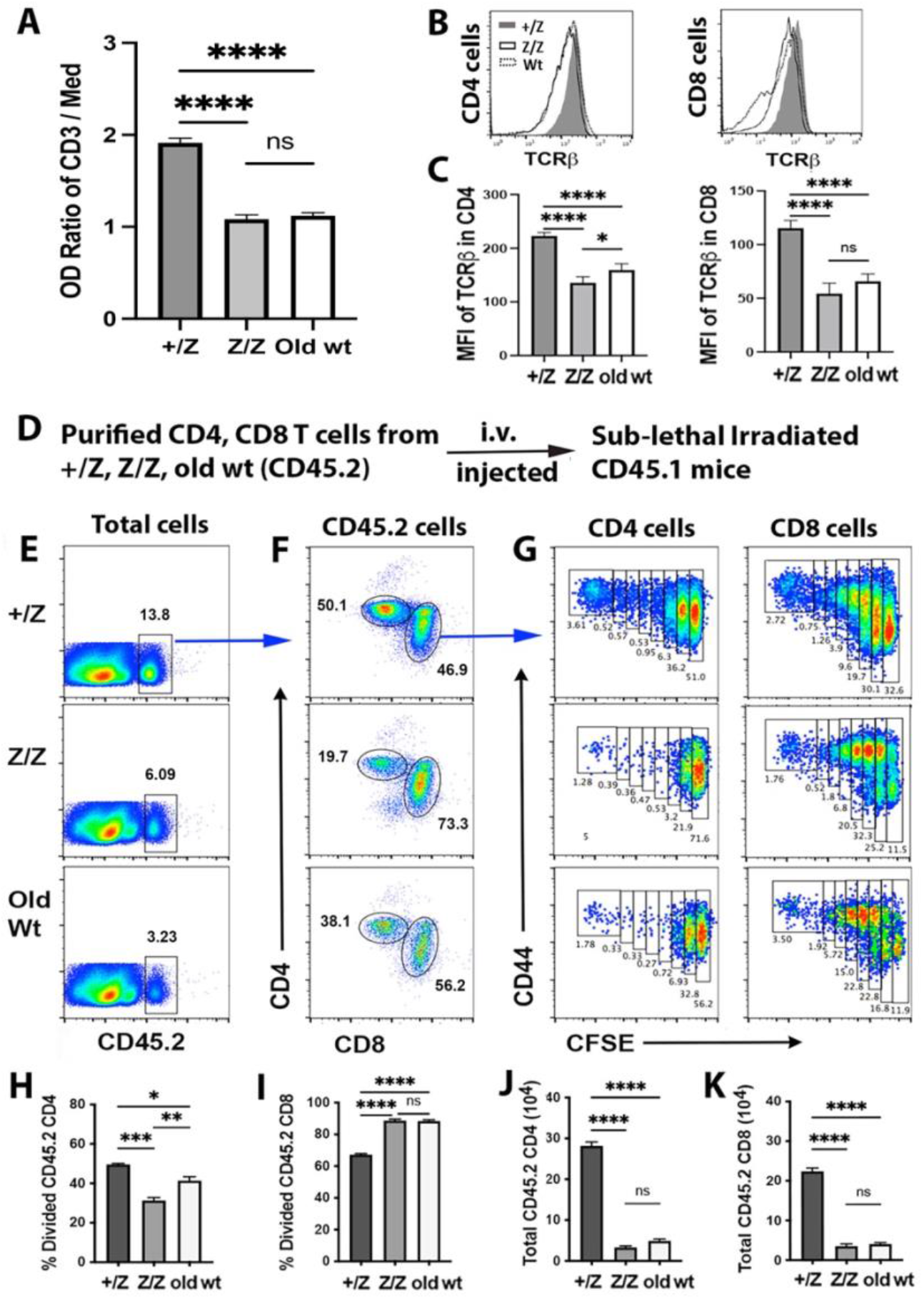
T cells from young adult *Foxn1^lacz^* mutants and aged mice had reduced proliferative capability after TCR stimulation. **(A).** Purified CD4 T cells from +/Z (3-months-old), Z/Z (3-months-old) and old wt (18-months-old) were co-cultured with or without anti-CD3 + CD28 antibodies in vitro for 48 hours and tOD values were measured. The OD ratio of anti-CD3 to medium is shown. **(B).** Histogram of TCRβ expression on gated CD4 (left) and CD8 (right) T cell. **(C).** TCRβ mean fluorescence density (MFI) on CD4 T cells (left) and **(C).** CD8 T cells (right). **(D).** Description of transfer experiment: 6 x 10^6^ cells/mouse of B cell-deleted CD4 and CD8 T cells isolated from +/Z (3-months-old), Z/Z (3-months-old), and wt (18-months-old) mice (CD45.2) were transferred into sub-lethally irradiated CD45.1 (2-3 mons) mice. Six days later, donor cells (CD45.2^+^) were analyzed. **(E).** Expression of CD45.2 on total spleen cells. **(F).** Expression of CD4 and CD8 on gated CD45.2. **(G).** The profiles of CFSE and CD44 on gated CD4 (left) and CD8 (right) T cells (CD45.2). **(H-I).** Percentage of cells with at least one division in CD45.2^+^ CD4 (H) and CD8 (I) T cells. **(J-K).** Total number of recovered CD45.2^+^ CD4 (J) and CD8 (K) T cells from spleen. (A): Two-way ANOVA; (C, H-K): One-way ANOVA. P values: ****≤0.0001, ***≤0.001, **≤0.01, *≤0.05.

To test the proliferative capability of T cells *in vivo*, CD4 and CD8 T cells from +/Z, Z/Z, and old wt mice were isolated and transferred into sub-lethally irradiated CD45.1 wt mice (Fig. 3.8 D). Six days later, donor T cells (CD45.2) were analyzed in host (CD45.1) mice. There was an increased frequency of CFSE^hi^ cells CD4 T cells in Z/Z and old wt compared to +/Z control, but cell division was reduced (Fig. 3.8 E-G, H). In contrast, there was a decreased frequency of CFSE^hi^ CD8 T cells in Z/Z and old wt compared to +/Z control while cell division was increased (Fig. 3.8 E-G, I). The absolute number of recovered CD45.2 donor cells was much less in both young Z/Z and old wt mice than in +/Z control (Fig. 3.8 J, K). These results indicate that peripheral CD4 and CD8 T cells from mice with premature or age-related thymic involution displayed weak TCR-CD3 stimulation that led to reduced activation and impaired homeostatic-driven proliferation *in vivo*.

### T cell cytokine production is similar between young *Foxn1^lacz^* and aged wild type mice

To test T cell function after activation, peripheral CD4 and CD8 T cell cytokine production was analyzed 5 and 24 hours after activation with aCD3+CD28 mAbs *in vitro*. Consistent with weak TCR activation, IL2 production was greatly reduced in CD4 T cells from old wt mice and slightly reduced in CD4 T cells from young Z/Z mice after 5 hours (Fig. 3.9 A, S-Fig. 3.5 A, B). After 24 hours, IL2 production was substantially reduced in CD4 T cells from both young Z/Z and old wt mice (Fig. 3.9 A, B). IFNγ levels were higher in CD4 T cells from old wt mice compared to young +/Z and Z/Z mice after 5 hours, but no significant differences were found after 24 hours (Fig. 3.9 A, C). CD8 T cells from both young Z/Z mutants and old wt mice had increased IFNγ production after 5 hours compared to +/Z. However, after 24 hours, IFNγ levels were similar between +/Z and Z/Z, and both were increased compared to old wt (Fig. 3.9 D and E). Together, these results suggest that CD4 T cells from young Z/Z mutants and old wt mice have impaired IL2 production, but both CD4 and CD8 cells from these mice can quickly produce the inflammatory cytokine IFNg in response to TCR signals.

**Figure 3.9.**
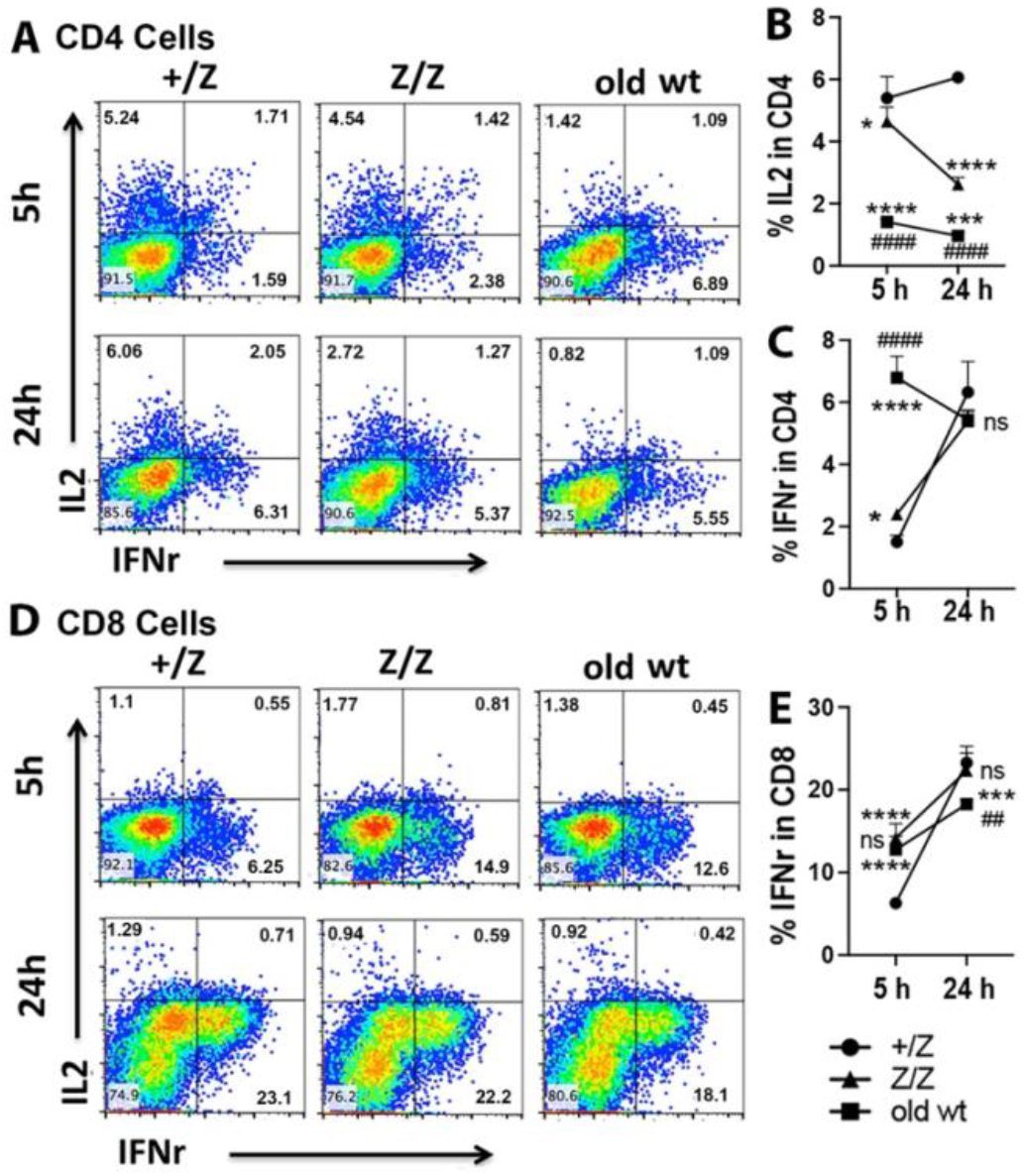
T cells from young adult *Foxn1^lacz^* mutants and aged mice had decreased production of cytokines after TCR-CD3 stimulation. 2x 10^6^ cells/well of B cell-deleted CD4 and CD8 T cells isolated from +/Z (3-months-old), Z/Z (3-months-old), and wt (18-months-old) mice (CD45.2) were co-cultured with or without anti-CD3 + CD28 in vitro for 5 hours (h) or 24 h. Cytokine production was analyzed for collected cells by intracellular staining. **(A).** Production of IL2 and IFNγ in CD4 T cells at 5 h (top) and 24 h (bottom). **(B).** Percentage of IL2 production in CD4 T cells at 5 h and 24 h. **(C).** Percentage of IFNγ production in CD4 T cells at 5 h and 24 h. **(D).** Production of IL2 and IFNγ in CD8 T cells at 5 h (top) and 24 h (bottom). **(E).** Percentages of IFNγ production in CD8 T cells at 5 h and 24 h. (B, C, and E): Two-way ANOVA. #: comparisons between +/Z and Z/Z or aged wt. *: comparisons between Z/Z and aged wt. P values: ****≤0.0001, ####≤0.0001, ***≤0.001, **≤0.01, ##≤0.01, *≤0.05. ns: no significance.

### The anti-influenza CD8 T cell response in the spleen is increased in young *Foxn1^lacz^* mutants compared to aged wild type mice

To investigate the effects of thymic involution on T cell immunity *in vivo*, we infected young +/Z, Z/Z, and old (16-months-old) wt mice with influenza and assessed morbidity (weight loss post-infection) and anti-viral T cell immunity. Overall, weight loss and cytokine production of CD4 T cells restimulated with HA peptide was similar (not significant) between all 3 groups except for a significant reduction of IFN-γ production in the spleen of old wt mice compared to +/Z control (data not shown). Next, we assessed the overall frequency and number of activated CD8 T cells reactive to the immunodominant NP (NP-tet^+^) and producing the inflammatory cytokines IFN-g and TNF-a in the lung and spleen at 10 days post infection (dpi), which is the peak of CD8 T cell response in C57BL/6 mice (Flynn et al. 1998). We performed a similar analysis on lymphocytes extracted from bronchoalveolar lavage (BAL), but the variability in cell extraction inherent to the isolation procedure restricted our analysis to cell frequency.

The only statistically significant change in the frequency of activated CD44^hi^ NP-tet^+^ CD8 T cells was found in the spleen, where there was an increase in Z/Z mice but a decrease in old wt compared to +/Z control (Fig. 3.10 A and B). The total number of activated CD44^hi^ NP-tet^+^ CD8 T cells was similar in Z/Z mice but decreased in old wt mice compared to control. A significant reduction of IFN-γ production in the spleen was only observed in old wt compared to +/Z controls (Fig. 3.10 C). In the lung, there was a trend of decreased CD44^hi^ NP-tet^+^ CD8 T cells and IFN-γ production in Z/Z and old wt. From BAL, the frequency of CD44^hi^ NP-tet^+^ CD8 T cells was slightly decreased in Z/Z, but no changes were found in IFN-γ production. Finally, there were no differences in TNF-a production between the three groups in any of the analyzed tissues (data not shown). Overall, these data demonstrate a similar CD8 T cell response to influenza in the respiratory tract of +/Z control mice, Z/Z mutants, and old wt mice. Interestingly, splenic T cells were significantly affected in old wt mice, and moderately affected in Z/Z mutants, suggesting that different subsets of responsive CD8 T cells were differentially affected by thymic involution.

**Figure 3.10.**
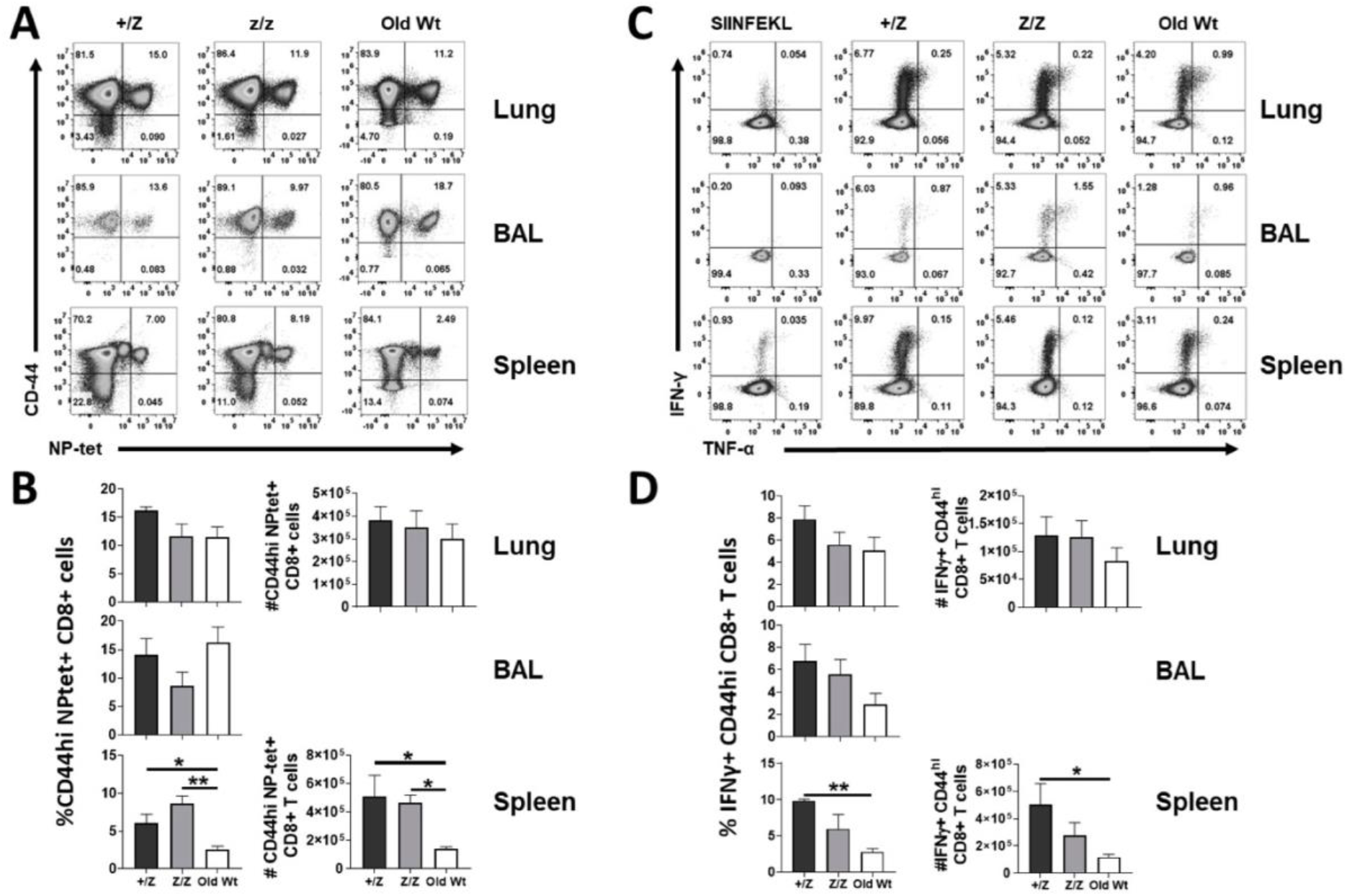
Premature involution in *Foxn1^lacz^* mice caused impaired response and function of activated antigen specific T cells after infection with influenza. +/Z, Z/Z, and old wt (16-months-old) mice were infected intranasally with 10^3^ pfu of influenza (x31). Ten days later, the overall frequency and number of activated CD8 T cells that were NP-tetramer positive or producing IFN-γ and TNF-a were measured. Representative flow plots (A) and their corresponding frequency and cell number of NP-tet+CD44+ cells of total CD8+ T cells from the lung, BAL, and spleen (B). Representative flow plots of IFN-γ^+^ and/or TNF-α^+^ cells (C) and the corresponding IFN-γ^+^ cells of CD8+CD44+ T cells from the lung, BAL, and spleen (D). Data is representative of 3 independent experiments of at least N=3 mice per group. Statistics according to one-way ANOVA using Graph Pad Prism 8 software. * = p≤0.05, ** = p≤0.01. Abbreviations: *BAL*, bronchoalveolar lavage.

## Discussion

Establishing a direct impact of thymic involution on the peripheral immune niches has been challenging due to various components contributing to the ageing peripheral immune system. The results of this study show an independent effect of thymic involution on the peripheral immune system distinct from the aging of hematopoietic stem cells and secondary lymphoid organs. Using single-cell RNA sequencing, we provided evidence of aged thymic microenvironments, especially in TECs from 1-month-old Z/Z compared to 18-month-old wt, which further supports the previously reported *Foxn1^lacz^* model (Chen et al. 2009) as a genetic model of premature onset and accelerated rate of thymic involution. We identified new stress-responsive cTECs and previously reported aging-associated TECs that lose epithelial markers whilst acquiring mesenchymal markers, potentially undergoing EMT. Notably, aaTECs that are exclusive to 18-month-old wt samples (Kousa et al. 2023) were present in 1-month-old Z/Z but were absent in 2-month-old wt samples, further supporting that deletion of *Foxn1* is sufficient to cause premature thymic involution.

As expected, thymopoiesis was affected by premature thymic involution in young *Foxn1^lacz^* including reduced thymic positive selection, increase thymic negative selection, and accelerated thymocyte maturation. Although there were some differences between young Z/Z and old wt mice, their changing trends were similar in both mice. Young Z/Z mice also exhibited a similar thymocyte phenotype to old wt (Table 1). Although there were some differences in proliferative and apoptotic thymocyte populations between young Z/Z and old wt compared to young +/Z control, young Z/Z mice overall displayed aged-like thymopoiesis similar to old wt. In young Z/Z, there was an overall increase in DN proliferation, which supports that thymic size reduction is caused by a decrease in *Foxn1* and is stromal-intrinsic. Thymocytes exhibited increased proliferative capacity in young Z/Z thymus compared to old wild-type thymus, suggesting relatively younger hematopoietic stem cells that are capable of homeostatic-driven proliferation. The most striking difference in young Z/Z was the significant decrease in CD4+ SP thymocytes compared to young +/Z and old wt, which could be due to the previously reported decrease in MHCII^hi^ and UEA-1^+^ cells and decreased proliferation of MHCII^lo^ TECs in young Z/Z mice (Chen et al. 2009). Alternatively, given the decrease in MHCII^hi^ TECs in naturally aging mice (Gray et al. 2006; Aw et al. 2008), different mTEC subset changes in young Z/Z compared to *de novo* mTEC aging may have caused the disparity. Increased negative selection in young Z/Z compared to young +/Z and old wt supports different mTEC subset compositions in young Z/Z.

**Table 3.1.**
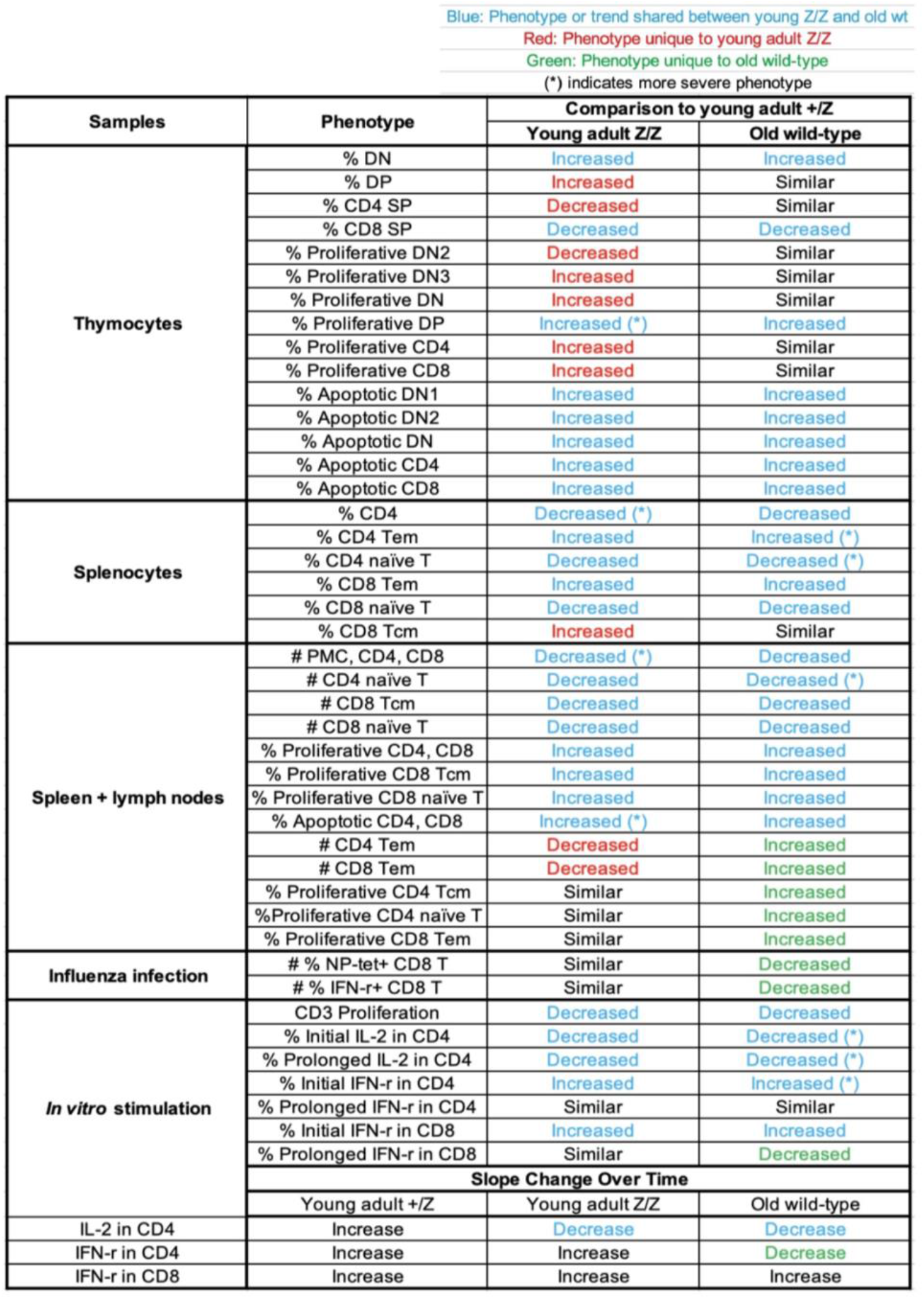
Summary of the T-cell phenotypes in young adult Z/Z and old wt.

scRNAseq of total splenocytes did not reveal significant differences between 1-month-old +/+ and Z/Z. The major differences were observed in 18-month-old wt, further demonstrating antigen exposures, physiological ageing, and HSC ageing shape the peripheral adaptive immune environment. Intriguingly, γδ T-cells are substantially decreased both in young Z/Z and old wt. There seems to be a direct relationship between the level of *Foxn1* and γδ T-cell production, which requires further investigation.

Decreased thymic output caused by premature thymic involution resulted in aged-like peripheral immune niches (Table 1). Reflecting a significant decrease in CD4 SP T-cells in the thymus, CD4+ cells in the spleen decreased significantly in young Z/Z. Further splenic T-cell analyses showed a decrease in naïve T-cell population with increased memory T-cell phenotypes in young Z/Z, closely resembling old wt. Moreover, we showed how premature thymic involution in *Foxn1^lacz^* mice resulted in lymphopenia, similar to *de novo* aged peripheral immune system. Due to prematurely decreased naïve T-cell production in the thymus, the absolute number of PMC, CD4, and CD8 were significantly decreased in Z/Z compared to +/Z control and old wt. With the decrease in naïve T-cell populations and increased T-cell proliferation and apoptosis, the peripheral immune profile of young Z/Z closely resembled that of old wt. However, we observed a huge disparity in the absolute number of CD4 and CD8 effector memory T-cell populations. These T cell populations were significantly decreased in young Z/Z, but were significantly increased in old wt compared to young +/Z control. Since the relative proportions of memory T-cells in young Z/Z is high akin to old wt, this disparity is likely caused by a more severe lymphopenia in young adult Z/Z with prematurely decreased naïve T-cell export from the thymus. We also observed a major difference in apoptosis between young Z/Z and old wt. While overall proliferation was similar between young Z/Z and old wt, apoptosis of CD4+ & CD8+ T-cells was significantly increased in young Z/Z compared to old wt and young +/Z control, further contributing to lymphopenia. Our data suggests overall peripheral immune niches of young Z/Z resembled those of old wt, but some differences existed due to severe lymphopenia in young Z/Z and potential antigen exposure in old wt.

Peripheral T-cell function in young Z/Z was also similar to old wt due to premature thymic involution (Table 1). *In vitro,* CD4 T-cell proliferative capacity was reduced by 50% upon TCR stimulation. We also observed distinct cytokine profiles across T-cells from young +/Z, Z/Z, and old wt. IL-2 maintenance in CD4+ T-cells was impaired in both young Z/Z and old wt, suggesting immunosenescence. Compared to a strong initial IL-2 response that increased over time in +/Z control, CD4+ T-cells from Z/Z displayed a strong initial IL-2 response that rapidly decreased in the span of 19 hours. In comparison, old wt CD4+ T-cells had weak IL-2 secretion that decreased further with time. On the contrary, both CD4+ & CD8+ T-cells had stronger initial IFN-γ response in young Z/Z and old wt, suggesting inflammaging.

Unlike *in vitro* stimulation, we did not observe statistically significant differences in T-cell immune response to influenza challenge in young Z/Z. We observed no difference in CD4+ T-cell responses, but a decreasing trend in antigen-specific and antigen-responsive CD8+ T-cells in the lung and airways of young Z/Z and old wt. The similarity in total cell count accompanied by functional decline is concordant with a previous report that thymectomized mice exhibit decreased influenza-responsive CD8+ T-cells, similar to aged mice due to a decrease in diversity of CD8 T-cells (Yager et al. 2008). Collectively, our findings support a direct effect of premature thymic involution on thymocytes and peripheral T-cells, which may provide an insight into which components contribute to immunosenescence and inflammaging.

## Acknowledgements

This work was supported by the Georgia Genomics and Bioinformatics Core and Center for Tropical and Emerging Global Diseases Flow Cytometry Facility at the University of Georgia. We thank the NIH Tetramer Core Facility (contract number 75N93020D00005) for providing the influenza NP MHC class I tetramer [H-2D(b)/ASNENMETM].

## Material and Methods

### Mice

Male and female C57BL6/J mice at 2-3 months and old mice at 15-18 months of age were purchased from National Institute on Aging. CD45.1 C57BL6/J mice were purchased from Jackson Laboratory. *Foxn1^+/lacz^* (+/Z) and *Foxn1^lacz/lacz^*(Z/Z) mice were generated on a C57BL6/J background as described previously (Chen et al. 2009; Gordon et al. 2007). All mice were maintained in a specific pathogen-free facility at the University of Georgia. The experiment was approved by the Institutional Animal Care and Use Committee of the University of Georgia.

### Isolation of the total thymic stroma

The thymus from three 1-month-old ZZ homozygotes and wild-type littermates were collected and pooled per single-cell dataset. Connective tissues, excess blood vessels, and adipocytes were carefully removed in 4℃ sterile 1X DPBS. Clean thymuses were then transferred to digestion buffer (2% FBS/1640 RPMI (w/ 25mM HEPES), DNase I (20µg/µL), Collagenase/Dispase (1mg/mL)) to be thinly sliced into 1∼2mm fragments. The digestion solution containing the thymic fragments was then incubated at 37℃ for 1 hour. In between the incubation period, the solution was gently mixed up and down with transfer pipettes every 20 minutes, 100 times each. Post-digestion, the solution was passed through a 70µm cell strainer to remove undigested fragments. Cells were centrifuged at 300 x g, 4℃ for 10 minutes. Red blood cells were then removed with a 2-minute incubation in RBC Lysis Buffer (10X) (BioLegend 420301). The solution was centrifuged again at 300 x g, 4℃ for 10 minutes. Cells were then resuspended in 3mL MACS buffer (filtered 0.5% BSA/2mM EDTA/1X DPBS). As per MACS separation guidelines, cells underwent CD45 separation with CD45 microbeads, mouse (Miltenyi Biotec 130-052-301).

### 10X library preparation and sequencing

Stromal cell viability was determined using a hemocytometer with Trypan Blue staining. Libraries were generated with samples with > 85% cell viability. Approximately 10,000 thymic stromal cells/splenocytes per sample were loaded into a 10X genomics Chromium instrument using a Single Cell 3’ Reagent Kit (v3). Post cDNA QC, the final DNA library was sequenced on Illumina NextSeq2000 P3.

### Single-cell RNA sequencing data analysis

As per the 10X guideline, we used CellRanger (v7) to generate and pre-process FASTQ files. CellRanger generated filtered_feature_bc_matrix files underwent QC steps and were analyzed using Seurat (v.4.3.0) in R (v.4.2.1). Logistic regression test was used for the differentially expressed gene analysis.

### Flow cytometry

Freshly isolated thymocytes in suspension (1.0 × 10^6^) were used for each sample. Cells were blocked by anti-CD16/32 (Clone:93) antibody before staining. For tracing the kinetic phenotypic profile and counting the numbers of thymocyte subsets, anti-CD4 APC-Cy7 or anti-CD4 violet 427 (GK1.5), anti-CD8 PerCp or anti-CD8 violet 570 (53-6.7), anti-CD44 FITC or PE (IM7), anti-CD25 APC (3C7), Anti-CD19 PE-Cy7 (6D5), anti-NK1.1 PE (PK136), anti-CD69 APC, anti-TCRβ PE-Cy7, and Qa2-biotin following PE-cy7 avidin were used to make a 6 or 8 color staining. For analysis of thymic progenitor Lin^-^ DN1a,b T cells in the total DN1 subpopulations, phycoerythrin (PE) conjugated lineage markers anti-CD3 (145-2C11), CD4, CD8, CD11c (N418), CD19, Gr-1 (RB68-C5), TER-119 (TER-119), NK1.1 antibodies were mixed and combined with anti-CD25 PerCp and anti-CD44 PE-Cy7, anti-CD117 APC (2B8), and CD24 FITC (M1/69) antibodies. For phenotypical analysis of spleen cells, anti-CD4 APC-Cy7 or anti-CD4 violet 427, anti-CD8 PerCp or anti-CD8 violet 570, anti-CD44 FITC, anti-CD62l PE-Cy7, anti-CD69 APC, anti-CD25 PerCp, anti-CCR7 PE, anti-CD45.1-PE, and anti-CD45.2-APC antibodies were used to make a 6 or 8 color staining. All antibodies were purchased from BioLegend (San Diego, CA). For AnnexinV staining (BioLegend, San Diego, Cat. # 640916), we followed the manufacturer instructions after surface antibodies staining. For antigen-specific CD8 T cell analysis, influenza nucleoprotein (NP) MHC class I tetramer [H-2D(b)/ASNENMETM], anti-CD8a FITC (53-6.7), anti-CD4 PE (RM4-5) and anti-CD44 PE-Cy7 (IM7) were incubated with cells for 1 hour at room temperature. All antibodies were purchased from Tonbo Biosciences (San Diego, CA) and the NP tetramer was generously provide by the NIH Tetramer Core Facility at Emory University (Atlanta, GA). Phenotypical analysis was performed with a Cyan ADP Flow Cytometer (Beckman Coulter, Miami, FL). The data were analyzed by Flowjo^TM^ Software (Tree Star, Ashland, OR).

### Thymic epithelial cell sorting

For thymic epithelial cell (TEC) sorting, thymic lobes were cut into 1 mm3 pieces and gently washed in 2% FBS + RPMI1640 medium to partially remove thymocytes. The thymic pieces were then transferred to a digestion solution containing collagenase/dispose (Roche) at 1mg/ml and DNase I 20ng/ ml in 8ml of 2% FBS + PRMI1640 medium, and then placed into a 37 C° water bath to digest for 60 minutes. The thymic tissue was gently agitated by plastic pipette every 20 minutes. The cells were filtered by passing through a size 70μm cell strainer and then incubated with anti-CD45 APC (30-F11), anti-EpCAM PE (G8.8) and anti-MHCII FITC (M5/114.15.2). TECs were sorted as CD45-EpCAM+MHCII+ cells by MoFlo^TM^ cell sorter.

### Cell culture and intracellular staining of cytokines

Peripheral lymphocytes were prepared from spleen and LNs of 2-3 months of age +/Z, Z/Z mice and 15-18 months of age BL6 wt mice. The cells (2.0 × 106/well) were plated in 24-well plates previously coated with 10 μg/ml anti-CD3ε (clone 145-2C11) and cultured in the presence of anti-CD28 at 2μg/ml for 5 or 24 h. As a control, cells were cultured in medium alone. To inhibit the secretion of newly synthesized cytokines, stimulations were conducted in the presence of 5 μg/ml brefeldin A (Sigma-Aldrich) in the last 5 hours of culture. After stimulation, cells were harvested and surface marker staining was performed with anti-CD4, anti-CD8, anti-CD44, CD25, and CD69. The cells were fixed in 0.5 ml/tube of fixation buffer (Cat. No. 420801) and permeabilized with permeabilization wash buffer (Cat. No. 421002) and then stained with anti-IL2 FITC, anti-IFNγ PE. Isotype of antibodies for each color was used as negative control. Stained cells were washed twice with permeabilization wash buffer and kept in FACS stock solution for flow cytometry analysis. Foxp3 staining was performed according to the protocol of the Treg staining kit from eBioscience. T cells from influenza infection experiments were re-stimulated after pulsing tissue-resident antigen-presenting cells (APCs) in the isolated tissues with either NP [ASNENMETM] or HA [YVQASGRVTVSTRRS] peptides, or a non-specific peptide [SIINFEKL] in RPMI supplemented with 10% FBS, 10% supplementum completum (HEPES, pen/strep, L-glutamine, β-mercaptoethanol, gentamycin, FCS, and RPMI) and golgistop for 5 hours at 37°C. Splenocytes from naïve mice were added to the airway extracted cells as an exogenous APC source. Subsequently, cells were surfaced stained with anti-CD8a FITC and anti-CD44 PE-CY7 for 20 minutes at 4°C and fixed overnight in 2% paraformaldehyde. The following day, cells were permeabilized using eBioscience PermWash (San Diego, California) for 30 minutes at 4°C, stained with anti-IFN-g APC (XMG1.2) and anti-TNF-a PerCP efluor 710 (MP6-XT22) for 20 minutes 4°C, and fixed in 2% paraformaldehyde for flow cytometry analysis. All antibodies purchased from Tonbo Biosciences (San Diego, CA) or eBioscience (Carlsbad, CA).

### Cell culture and proliferation

Peripheral lymphocytes were isolated from the spleen and lymph nodes of 2-3 months-old +/Z and Z/Z mice and 15-18 months-old BL6 wild-type mice. CD4+ T cells were purified by incubation with anti-CD4 microbeads and passing through a magnetic column (Miltenyi Biotec). The purity of cells was higher than 95% as determined by flow cytometry. Purified CD4+ cells (2.0 × 105/well) were then placed in 96-well plates previously coated with 10 μg/ml anti-CD3ε Ab. Anti-CD28 was added at 2 μg/ml. Cells were cultured in a 37°C, 5% CO2 incubator for 48 hours. For the last 4 hours, cultures were pulsed with CellTiter 96 Aqueous One Solution Reagent (Promega), 20 μl/well. As a control, cells were cultured in medium alone. We measured and recorded the absorbance at 490 nm using a 96-well plate reader. Some cells were also collected for phenotypic analysis with anti-CD4, CD8, CD25, CD69 antibodies.

### Chimera generation

For analysis of peripheral lymphopenia, purified CD4 and CD8 T cells isolated from CD45.1 mice were transferred into Foxn1 Z/Z and old wt mice. For analysis of the proliferative capability of T cells, purified CD4 and CD8 T cells isolated from Foxn1 Z/Z and old wt (CD45.2) mice were transferred into sublethal irradiated (500 Rad) CD45.1 wt mice. CD4 and CD8 T cells isolated from spleen and lymph nodes were purified by incubating with anti-CD4 and CD8 microbeads (Miltenyi Biotec) and passing through LS magnetic columns following manufacturer instructions. The purity of cells was higher than 95%. Purified T cells were then incubated with CFSE (Molecular Probes) at final concentrate of 10 mM in PBS + 0.1% BSA, 37°C, for 10 min. The unlabeled CFSE was quenched by adding 10% FCS medium and then washed by 2% FBS medium. Depending on the experiment, Foxn1 Z/Z and old wt or sublethal irradiated BL6 CD45.1 wt mice were retro-orbitally injected with CFSE-labeled cells (4.0 x 10^6^/mouse). Six days after the transfer, peripheral lymphocytes collected from host lymph nodes and spleen were stained and analyzed by flow cytometry.

### BrdU incorporation and staining

Mice were injected once intraperitoneally with 1 mg of 5-bromo-2’-deoxyuridine (BrdU, Sigma-Aldrich) and then fed with BrdU-containing water (0.8 mg/ml) for five days. The thymocytes were harvested and stained with CD4, CD8, CD25, CD44 antibodies. The surface-stained cells were fixed and permeabilized in PBS containing 1% paraformaldehyde plus 0.01% Tween20 for 48 hours at 4°C and then submitted to the BrdU DNase1 detection technique. FITC-anti-BrdU Ab (clone 3D4; BioLegend) was used for BrdU staining.

### Influenza infection of mice and single cell suspension preparation

All mice were intranasally infected with 10^3^ pfu of influenza A virus A/HK-x31 (x31, H3N2) in 50 ul of PBS. At 10 days post infection (dpi), mice were sacrificed to obtain single cell suspensions from the lung tissue, lung airways, and spleen. Briefly, an incision was made into the trachea to perform a bronchoalveolar lavage (BAL) by dispensing 1 mL of PBS into the lung and aspirating the same volume. Lungs were then perfused with PBS/heparin, excised, and collected into 5 mL RPMI. Lastly, spleens were excised and collected into 5 mL RPMI. For tissue digestion, spleens were mechanically disrupted and passed through 40 uM nitex nylon mesh. Red blood cells were removed by incubation with tris ammonium chloride for 2 minutes at room temperature. Lungs were minced before incubation for 30 minutes at 37°C with 1.25 mM EDTA followed by incubation in 150 units/mL collagenase. Lungs were then passed through a 40 µM cell strainer to obtain cells, resuspended in 44% Percoll, and underlaid with a 67% Percoll solution. After centrifugation at 2800 rpm for 20 minutes, lymphocytes were then collected at the cellular interface.

### Statistical analysis

All data were collected in a Microsoft Excel file and analyzed using Prism 8 software by one-way or two-way analysis of variance (ANOVA).

