## Supplementary figures for "Premature thymic involution in young *Foxn1^lacz^* mutant mice causes peripheral T cell phenotypes similar to aging-induced immunosenescence"

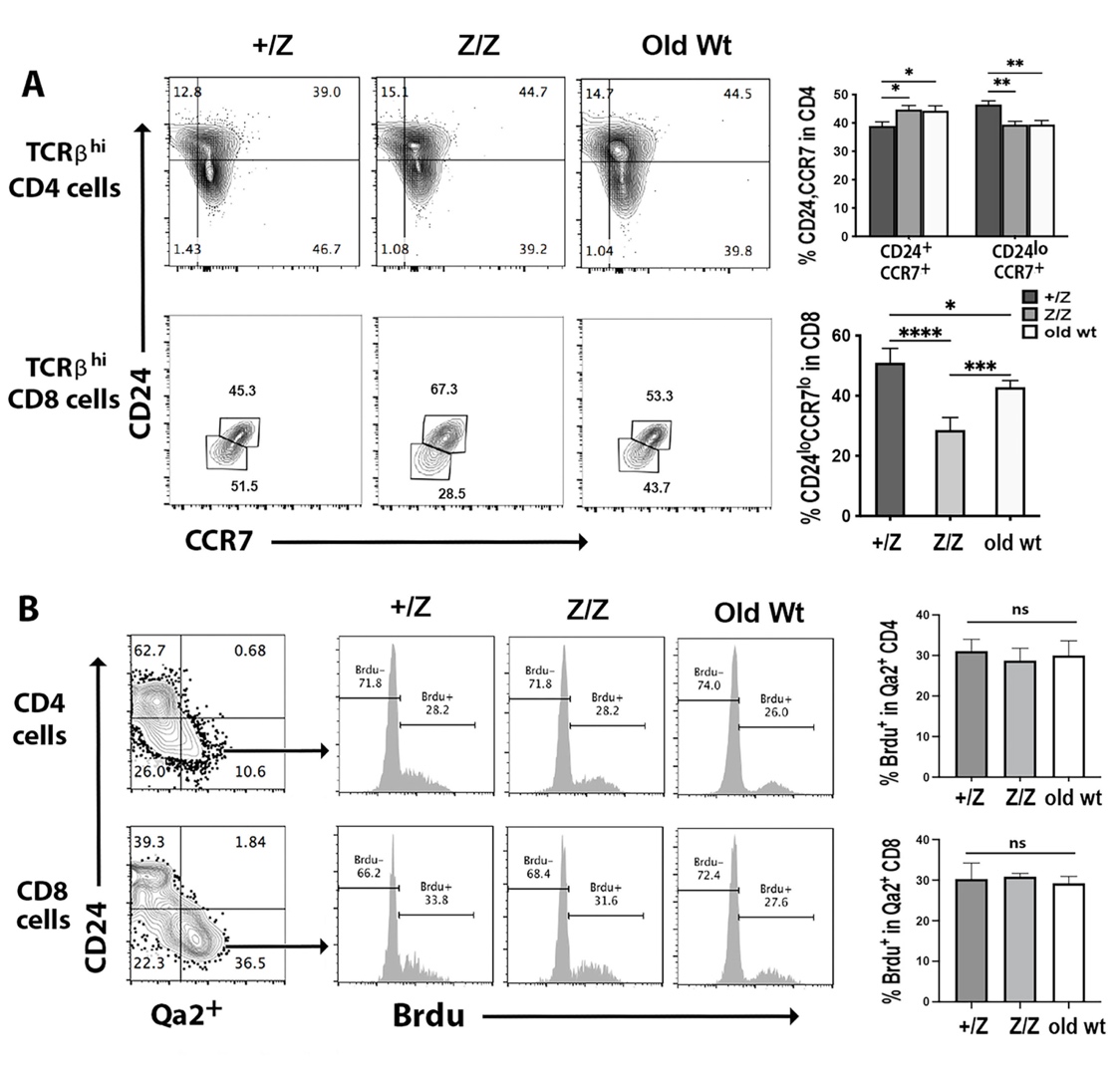

**Supplementary Figure 3.1. Down-regulation of CD24 was reduced in young *Foxn1^lacz^* mutants and old wt mice.**

Flow cytometric analysis of thymocytes from +/Z (2-months-old), Z/Z (2-months-old) and old wt mice (18-months-old). **(A).** Expression of CD24 and CCR7 on TCRβ^hi^ CD4 and CD8 SP thymocytes. **(B).** Expression of CD24 and Qa2 on gated CD4 (top) and CD8 (bottom) SP thymocytes (left). Histogram of Brdu staining on TCRβ^+^ Qa2^+^CD4 and TCRβ^+^ Qa2^+^CD8 SP thymocytes (middle). Summary data of percentage of Brdu^+^ in Qa2^+^ CD4 and CD8 SP thymocytes on the right. Data are representative of three individual experiments with at least 5 mice per group.

(A for CD4 cells): Two-way ANOVA; (A for CD8 cell and B): One-way ANOVA. P values: ****≤0.0001, ***≤0.001, **≤0.01, *≤0.05.

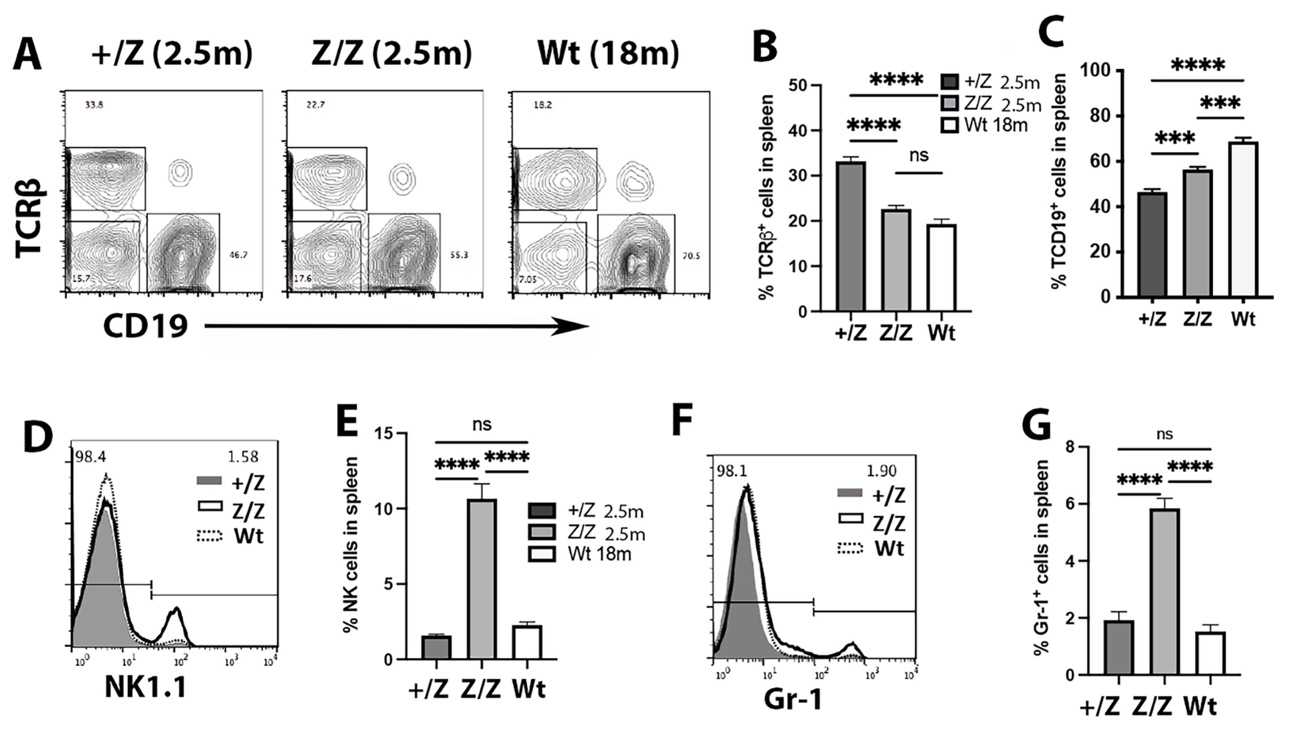

**Supplementary Figure 3.2. Percentage of splenic B cells, NK cells, and granulocytes were increased in 2.5-months-old *Foxn1^lacz^* mutants.**

Flow cytometric analysis of splenocytes from +/Z (2.5-months-old), Z/Z (2.5-months-old) and aged wt mice (18-months-old). **(A)** Histogram of TCRβ and CD19 total spleen cell expression. **(B).** Percentage of TCRβ^+^ spleen cells. **(C).** Percentage of CD19^+^ spleen cells. **(D).** Histograms of NK1.1-expressing spleen cells. **(E).** Percentage of natural killer (NK) cells in the spleen. **(D).** Histogram of Gr-1-expressing spleen cells. **(D).** Percentage of granulocyte in the spleen. Data are representative of three individual experiments with at least 5 mice per group. (B, C, E and G): One-way ANOVA. P values: ****≤0.0001, ***≤0.001, ns: No significance. Abbreviations: *m,* months.

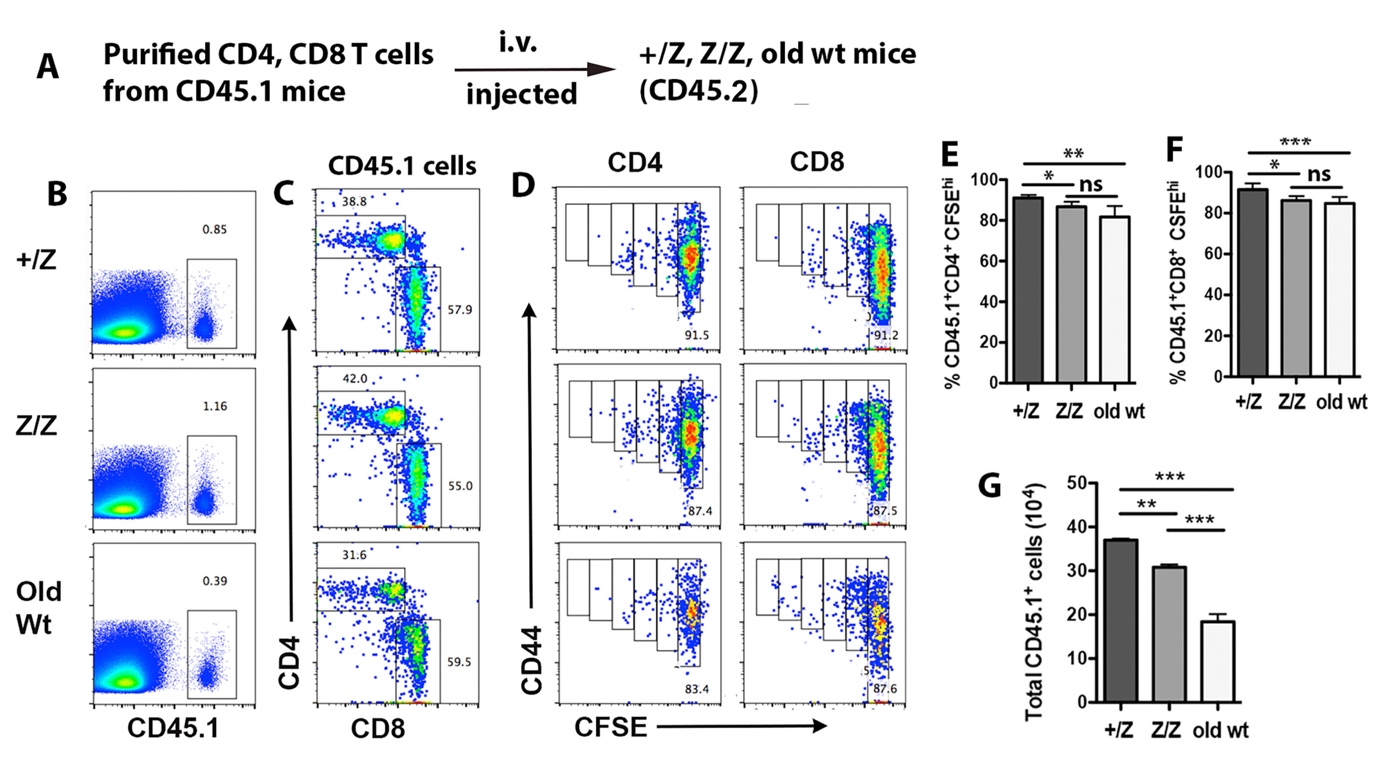

**Supplementary Figure 3.3. Peripheral lymphopenia promoted weak homeostatic proliferation in young *Foxn1^lacz^* mutants and aged wild type mice.**

6 x 10^6^ cells/mouse of B-cell-deleted CD4 and CD8 T cells isolated from CD45.1 mice were transferred into +/Z (2-3 months-old), Z/Z (2-3 months-old) and old wt (18-months-old) mice (CD45.2). After 6 days, the donor cells (CD45.1^+^) were analyzed from hosts by showing the gate of CD45.1 cells. **(A).** Description of cell transfer experiment. **(B).** CD45.1 expression on total spleen cells. **(C).** CD4 and CD8 expression on gated CD45.1 positive cells. **(D)** Profile of CFSE and CD44 expression on gated donor CD4 and CD8 T cells. **(E).** Histogram of CFSE^hi^ percentage in CD45.1^+^ CD4 T cells. **(F).** Histogram of CFSE^hi^ percentage in CD45.1^+^ CD8 T cells. **(G).** Total number of donor lymphocytes (CD45.1) recovered from each host. Data are representative of two individual experiments with at least 5 mice per group. (E- G): One-way ANOVA. P values: ***≤0.001, **≤0.01, *≤0.05. ns: No significance.

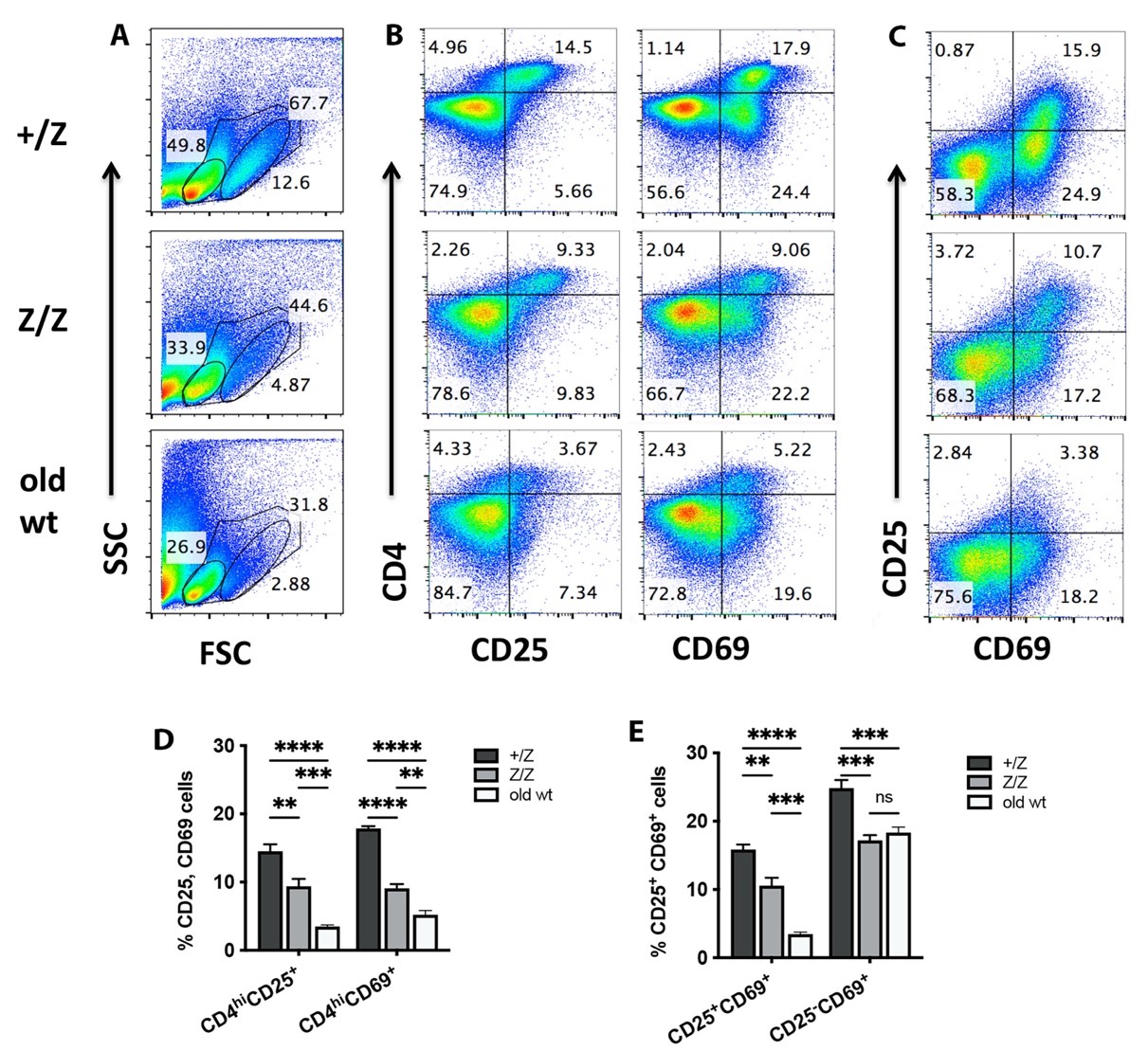

**Supplementary Figure 3.4. Peripheral CD4 T cells displayed weak activation upon stimulation of TCR-CD3 signaling *in vitro*.**

Purified CD4 T cells from +/Z (3-months-old), Z/Z (3-months-old) and old wt mice (18-months-old) were co-cultured with or without anti-CD3 + CD28 antibodies for 48 hours *in vitro*. Activated markers CD25 and CD69 were analyzed on collected cells by flow cytometry. **(A).** Profiles of FSC and SSC for cell ages. **(B).** Expression of CD25 vs CD4 (left) and CD69 vs CD4 (right) on gated live cells. **(C).** Expression of CD25 and CD69 on gated live cells. **(D).** Percentage of CD4^hi^CD25^+^ and CD4^hi^CD69^+^ cells in gated live cells. **(E).** Percentage of CD25^+^CD69^+^ and CD25^-^CD69^+^ cells

in gated live cells. Data are representative of two individual experiments with at least 5 mice per group. (D and E): One-way ANOVA. P values: ****≤0.0001, ***≤0.001, **≤0.01, ns: No significance.

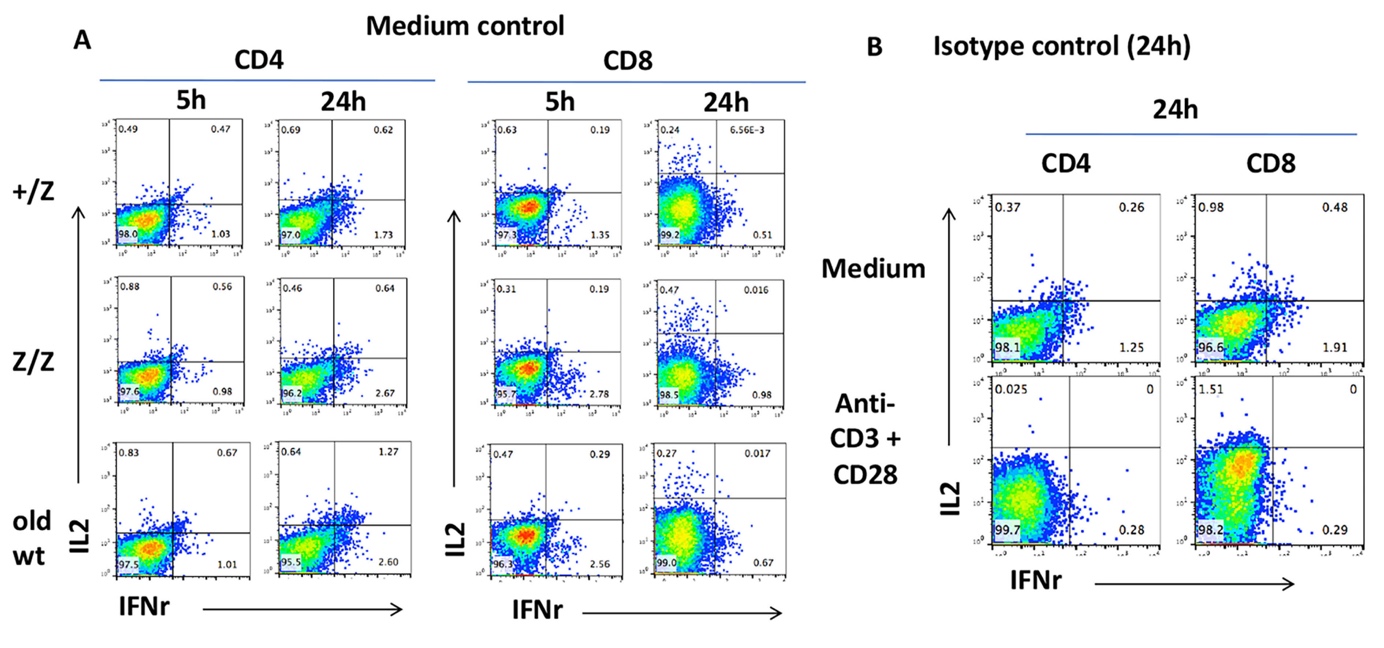

**Supplementary Figure 3.5. Intracellular cytokine staining of medium and isotype control.**

2x 10^6^ cells /well of B cell-deleted CD4 and CD8 T cells isolated from +/Z (3-months-old), Z/Z (3-months-old), and wt (18-months-old) mice (CD45.2) were co-cultured with anti-CD3 + CD28 and/or medium only *in vitro* for 5 hours (h) or 24 h. Cytokine production was analyzed for collected cells by intracellular staining. **(A).** Production of IL2 and IFNγ at 5 h and 24 h in CD4 T cells (left) and in CD8 T cells (right). **(B).** Profiles of isotype-FITC and isotype-PE reagents that matched IL-2-FITC and IFNγ-PE antibodies at 24 h post-activation of CD4 (left) and CD8 T cells (right) with medium (top) or in anti-CD3 + CD28 (bottom).
